# Retrovirus-derived *RTL9* plays an important role in innate antifungal immunity in the eutherian brain

**DOI:** 10.1101/2023.06.29.547130

**Authors:** Fumitoshi Ishino, Johbu Itoh, Masahito Irie, Ayumi Matsuzawa, Mie Naruse, Toru Suzuki, Yuichi Hiraoka, Tomoko Kaneko-Ishino

## Abstract

Retrotransposon Gag-like (RTL) genes plays a variety of essential/important roles in the eutherian placenta and brain. It has recently been demonstrated that *RTL5* and *RTL6* (aka *sushi-ichi retrotransposon homolog 8* (*SIRH8*) and *SIRH3*) are microglial genes that play important roles in the brain’s innate immunity against viruses and bacteria by their removal of double-stranded RNA and lipopolysaccharide, respectively. Here we demonstrate that *RTL9* (aka *SIRH10*) also plays an important role, degrading fungal zymosan in the brain. The RTL9 protein is localized in the microglial lysosomes where incorporated zymosan is digested. Interestingly, in *Rtl9* knockout mice expressing RTL9ΔC protein lacking the C-terminus retroviral GAG-like region, the zymosan degrading activity was lost, demonstrating that RTL9 is essentially engaged in this reaction, presumably via its GAG-like region. Together with our previous study, this result highlights the importance of three retrovirus-derived microglial RTL genes as eutherian-specific constituents of the current brain innate immune system, *RTL9*, *RTL5* and *RTL6* responding to fungi, viruses and bacteria, respectively.

**Author Summary:** We have recently demonstrated that *RTL5* and *RTL6* are microglial genes that play important roles in the brain’s innate immunity against viruses and bacteria. In this report, we demonstrate that *RTL9* is functional in innate antifungal immunity in the brain because *Rtl9* KO mice lose zymosan degradation activity. Fungi are one of the most dangerous infectious pathogens, along with viruses, bacteria and protozoa. Phagocytic cells, such as microglia/macrophages, are essential for mounting a defense against fungal infection. As defects in these cells reduce host resistance to fungal infection, *RTL9* is an antifungal therapy target as a newly identified member of innate antifungal immunity in eutherians.

## Introduction

Retrotransposon-Gag like (RTL, aka sushi-ichi retrotransposon homolog (SIRH)) genes are eutherian-specific except for therian-specific *Paternally expressed 10* (*PEG10*): comparative genome analyses have provided convincing evidence that *PEG10* emerged in a therian common ancestor while others are conserved exclusively in eutherians (Ono et al., 2001 and 2006; Brandt et al., 2005; Youngson et al., 2005; Suzuki et al., 2007; Edwards et al., 2008; Kaneko-Ishino and Ishino, 2012), indicating that they are presumably derived from an extinct retrovirus similar to the sushi-ichi retrotransposon (Imakawa et al., 2022). This view is supported by the evidence that the gypsy retrotransposon to which the sushi-ichi retrotransposon belongs is an infectious retrovirus of *Drosophila melanogaster*, possessing an *ENV*-like gene in addition to *GAG* and *POL* (Kim et al., 1994; Song et al., 1994).

Among the 11 RTL genes*, PEG10*, *RTL1* (aka *PEG11*) and *Leucine zipper down-regulated in cancer 1* (*LDOC1*, aka *SIRH7* and *RTL7*) play essential but different roles in the placenta (Ono et al., 2006; Sekita et al., 2008; Naruse et al., 2014; Kitazawa et al., 2017) while *RTL4* (aka *SIRH11* and *Zinc finger CCHC domain-containing 16* (*ZCCHC16*) plays an important role in controlling impulsivity in the brain (Irie et al., 2015) and is recognized as a causative gene in autism spectrum disorders (Lim et al., 2013). Moreover, *RTL1* plays important roles in the muscle and brain and is considered to be one of the major genes responsible for Kagami-Ogata and Temple syndromes (Kagami et al., 2008 and 2015; Ioannides et al., 2014; Kitazawa et al., 2020 and 2021; Chou et al., 2022).

We have recently demonstrated that *RTL5* (aka *SIRH8*) and *RTL6* (aka *SIRH3*) are microglial genes playing important roles in the brain innate immunity against viruses and bacteria by their removal of double stranded (ds)RNA and lipopolysaccharide (LPS), respectively (Irie et al., 2022). Microglia are the primary immune cells in the brain, playing a central role in the innate immune responses to various pathogens via a variety of Toll-like receptors (TLRs) that induce inflammation via regulation of cytokine and interferon expression (Hanisch and Kettenmann, 2007; Norris and Kipnis, 2018). It is well known that the TLR system is widely conserved in the animal kingdom from worms to mammals (Akira et al., 2006), while *RTL5* and *RTL6* are apparently novel constituents of the eutherian innate immune system.

Innate antifungal immunity is another important survival mechanism of organisms in addition to those against bacteria and viruses. Fungal diseases are associated with increased morbidity and mortality, particularly in immunocompromised individuals (Feldman et al., 2018; Brown, 2011). Zymosan is frequently used to experimentally induce an inflammatory reaction, as it mimics fungal infection. It is a component of the yeast cell wall that consists of protein-carbohydrate complexes, such as glucans, mannans, proteins, chitins and glycolipids. It is recognized by several cell surface receptors, such as TLR2, TLR6 and Dectin-1, becomes incorporated into phagosomes and is ultimately degraded in the phagolysosomes present in macrophages/microglia (Underhill et al. 1999; Underhill. 2003, Salazar and Brown. 2018; Fieldman et al., 2019).

Phagocytes, including macrophages/microglia, play a central role in antifungal immunity using a variety of oxidative and non-oxidative mechanisms that work synergistically to kill extracellular and internalized fungi by producing reactive oxygen intermediates and/or hydrogen peroxide, such as hypochlorous acid and hypoiodous acid in the former activity and antibacterial peptides and hydrolases in the latter (Brown, 2011). Lysosomes are organelles in animal cells that act as degradation centers for a variety of biomolecules, including carbohydrates, proteins, lipids and nucleotides, with various hydrolytic enzymes functioning in an acidic environment (pH4.5-5.0) (Luizo et al., 2007; Xu and Ren, 2015).

*RTL9* (aka *SIRH10* or *Retrotransposon GAG domain containing 1* (*RGAG1*)) is another RTL gene that is highly conserved in eutherians, but the biological function of which has remained undetermined. In this report, we show that *RTL9* localizes in microglial lysosomes and plays an essential role in the degradation of fungal zymosan. Thus, it turns out that eutherians came to possess at least three eutherian-specific pathways against fungi, viruses and bacteria as a result of the respective acquisition of *RTL9, RTL5* and *RTL6* in the innate immune system of the brain.

## Results

### *RTL9* is conserved in eutherians

*RTL9* encodes a large protein comprising 1347 amino acids (aa) and is evolutionarily conserved in the eutherians (dN/dS <1) (Table 1 and Fig. S1), suggesting it confers certain evolutionary advantage(s).

**Table 1.** Pairwise dN/dS ratio of RTL9 in 10 representative eutherian species. The human, chimpanzee, mouse, rat (Euarchontoglires), dog and horse (Laurasiatheria), manatee and elephant (Afrotheria) and armadillo and sloth (Xenarthra) species represent all four major groups of eutherians.

| dN/dS | human | chimpanzee | mouse | rat | dog | horse | manatee | elephant | armadillo |
| --- | --- | --- | --- | --- | --- | --- | --- | --- | --- |
| chimpanzee | 0.7009 |  |  |  |  |  |  |  |  |
| mouse | 0.4544 | 0.4529 |  |  |  |  |  |  |  |
| rat | 0.4022 | 0.4089 | 0.3364 |  |  |  |  |  |  |
| dog | 0.4272 | 0.4408 | 0.4236 | 0.3936 |  |  |  |  |  |
| horse | 0.6116 | 0.6153 | 0.4687 | 0.4153 | 0.4181 |  |  |  |  |
| manatee | 0.5264 | 0.5334 | 0.4170 | 0.4060 | 0.4148 | 0.5173 |  |  |  |
| elephant | 0.5409 | 0.5488 | 0.4181 | 0.4178 | 0.4751 | 0.5973 | 0.4284 |  |  |
| armadillo | 0.5552 | 0.5638 | 0.5075 | 0.4542 | 0.4794 | 0.5588 | 0.5227 | 0.5334 |  |
| sloth | 0.6619 | 0.6803 | 0.5726 | 0.5493 | 0.5014 | 0.6432 | 0.6154 | 0.6365 | 0.7614 |

Like the other RTL genes, *RTL9* exhibits homology to the sushi-ichi retrotransposon GAG, but is unique in that it further exhibits homology to two herpes virus-like sequences, the outer envelope glycoprotein BLLF1 (super family member pfam05109, the gray box in the figure corresponding to 280-594 aa) and the large tegument protein UL36 (super family member pha03247, the orange box corresponding to 787-1141 aa) (Fig. 1A and Figs. S1 and S2). The BLLF1 and UL36 proteins have several functional motifs and domains, but there are no such functional sequences present in RTL9 (Figs. S3 and S4, see also the Discussion section). More detailed homology analysis has indicated that sushi-ichi GAG in face exhibits a high degree of homology with a widespread region, including the capsid domain (the blue box corresponding to 1169-1250 aa) (Fig. 1A). One such region is the one corresponding to 1148-1367 aa, from just before the capsid region to the end of RTL9 (the minimal GAG-like region, the red dashed box in Fig. 1A, identity 25.5%, similarity 43.1%) (Fig. S5) with another being the region from 927 to 1367 aa (the maximal GAG-like region indicated by the red double-headed arrow in Fig. 1A, identity 21.8%, similarity 35.7%) corresponding to the entire GAG sequence (Fig. S6). Thus, it remains ambiguous whether the 927-1148 aa portion originated from UL36 or GAG. In any event, information from the protein sequence alone is not sufficient to infer the biological function of the RTL9 protein.

**Figure 1.**
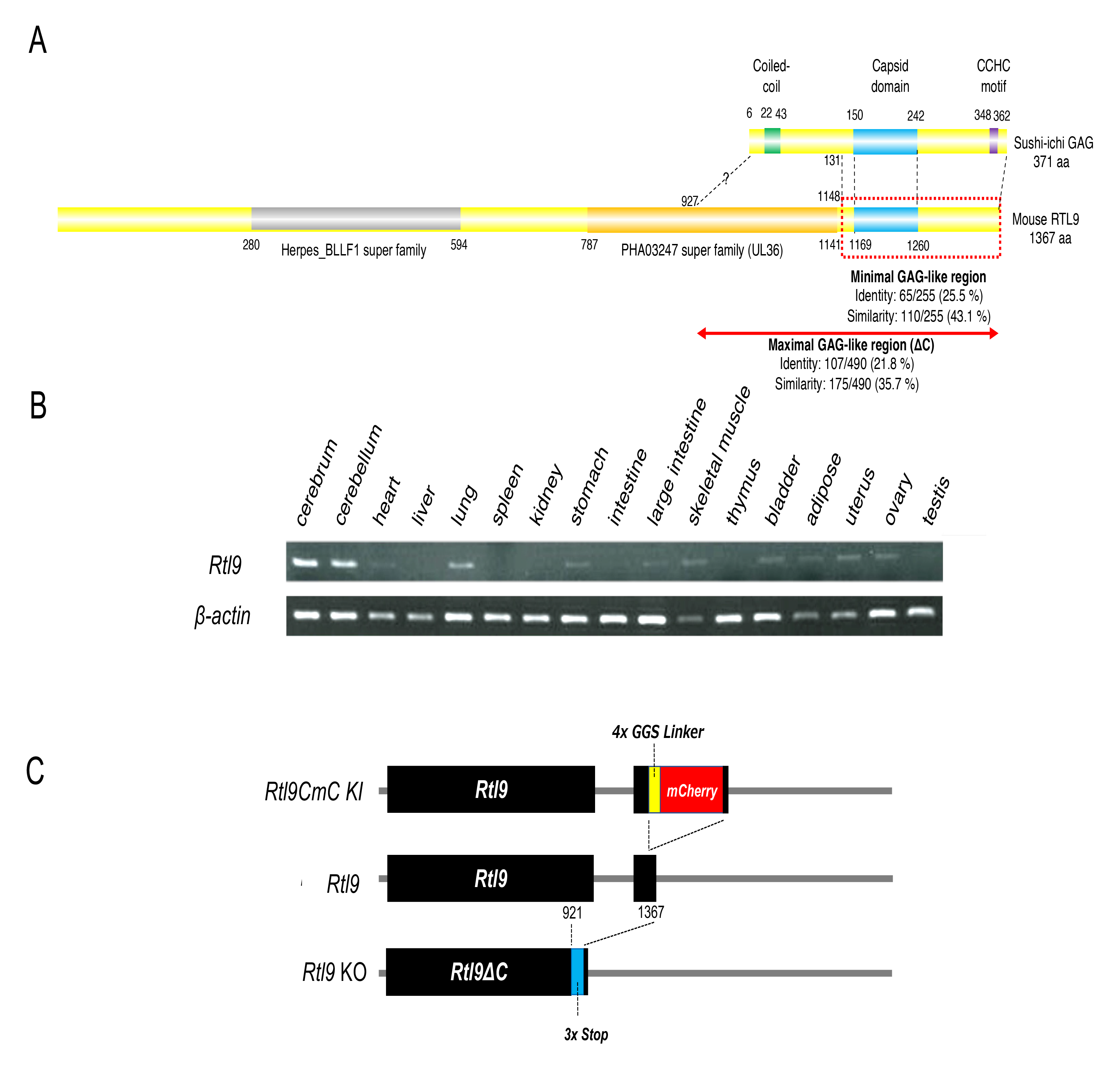
Features of the RTL9 protein, mRNA expression, and gene structures of KI, normal and KO mice. **A**) Alignment of the sushi-ichi GAG and mouse RTL9 protein. Coiled-coil motifs in the N-terminus, a capsid domain in the middle and a zinc finger CCHC domain in the C-terminus are represented as green, blue and purple boxes, respectively. The Herpes BLLF1 super family and PHA03247 super family (UL36) motifs are depicted with a gray and orange box, respectively, and the capsid domain of the sushi-ichi retrotransposon is presented as a blue box. Minimal and maximal Gag-like regions are presented with a red dashed line and indicated by a red double-headed arrow, respectively. **B)** Mouse *Rtl9* mRNA expression in the adult tissues and organs at 19 week. The RT-PCR products using total RNA (10 ug, 32 cycles for *Rtl9* and 26 cycles for *β-actin* as a control). **C)** The generation of the*Rtl9CmC* KI and *Rtl9* KO mice. Top: the open frame of the mCherry protein (red) is fused with the C-terminus of RTL9 via a 4 x GGS linker (yellow). Middle: mouse *Rtl9* comprising two exons. Bottom: the RTL9 C-terminal region is deleted by the insertion of a 3 x stop sequence (blue) at the 921 aa position.

*Rtl9* mRNA expression was very weak, detected only in the brain, lung, skeletal muscle and uterus after 32 PCR cycles, and was almost absent from most tissues and organs at 19 weeks (19w) in adult mice (Fig. 1B). Similar results were reported for human *RTL9* (GTEx Portal site, searched as *RGAG1*): low expression in the brain, heart and ovary in addition to a relatively higher expression in the testis (Fig. S7). Therefore, for the *in vivo* detection of RTL9, we generated an *Rtl9-mCherry* knock-in (KI) mouse that expresses the RTL9-mCherry (RTL9CmC) protein fused with a fluorescent mCherry protein after the C-terminus of endogenous RTL9 (Fig. 1C, top and Fig. S8). We also generated an *Rtl9ΔC* mouse (hereafter called the *Rtl9* KO mouse) that produces RTL9 with a truncated C-terminus (922-1367 aa, the maximal GAG-like region, Fig. 1A and Fig. S9) to elucidate the function of RTL9, especially of its C-terminal GAG-like region and potentially including the last portion of UL36-like region (927-1145 aa).

### RTL9 localizes in microglial lysosomes

The RTL9CmC protein is highly expressed in microglial lysosomes in the neonatal brain. Using *Rtl9-CmCherry* KI mice, we detected the mCherry signal in the midbrain and around the thalamus in postnatal day 0-15 (P0-15) neonates in regions such as the superior colliculus (SC), cerebral aqueduct (AQ), periaqueductal gray (PAG) matter, posterior commissure (PC) and the commissures of the superior and inferior colliculus (CSC and CIC) (Fig. 2A, left). The mCherry signal was specific to the *Rtl9-CmCherry* KI brain and almost negligible in the control WT brain (Fig. 2A, right), confirming that the mCherry signal correctly reflects the localization of the RTL9CmC protein. Its expression profile remained almost the same until 3∼4 weeks of age when the signal strength progressively diminished, and this trend continued up to the adult period.

**Figure 2.**
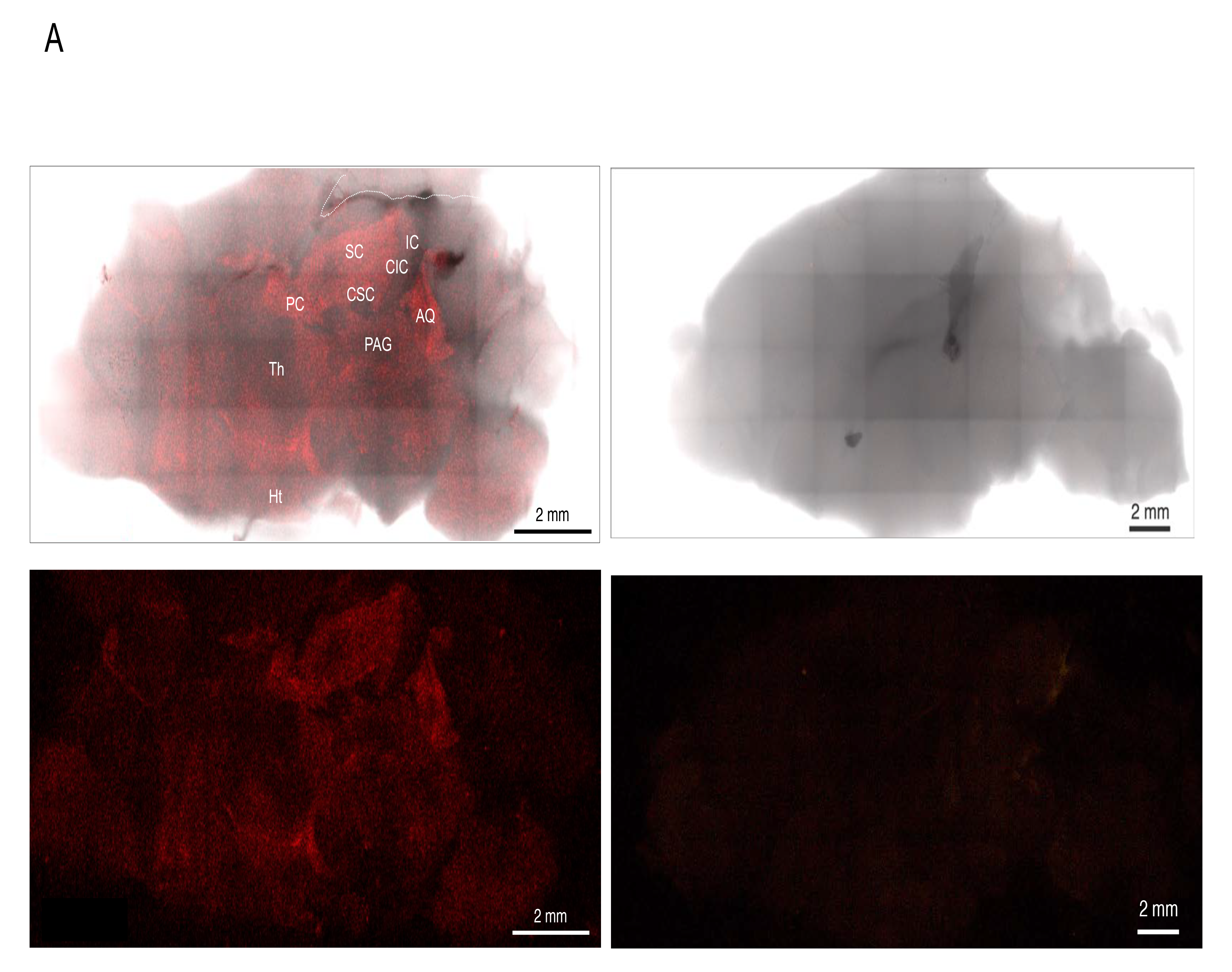
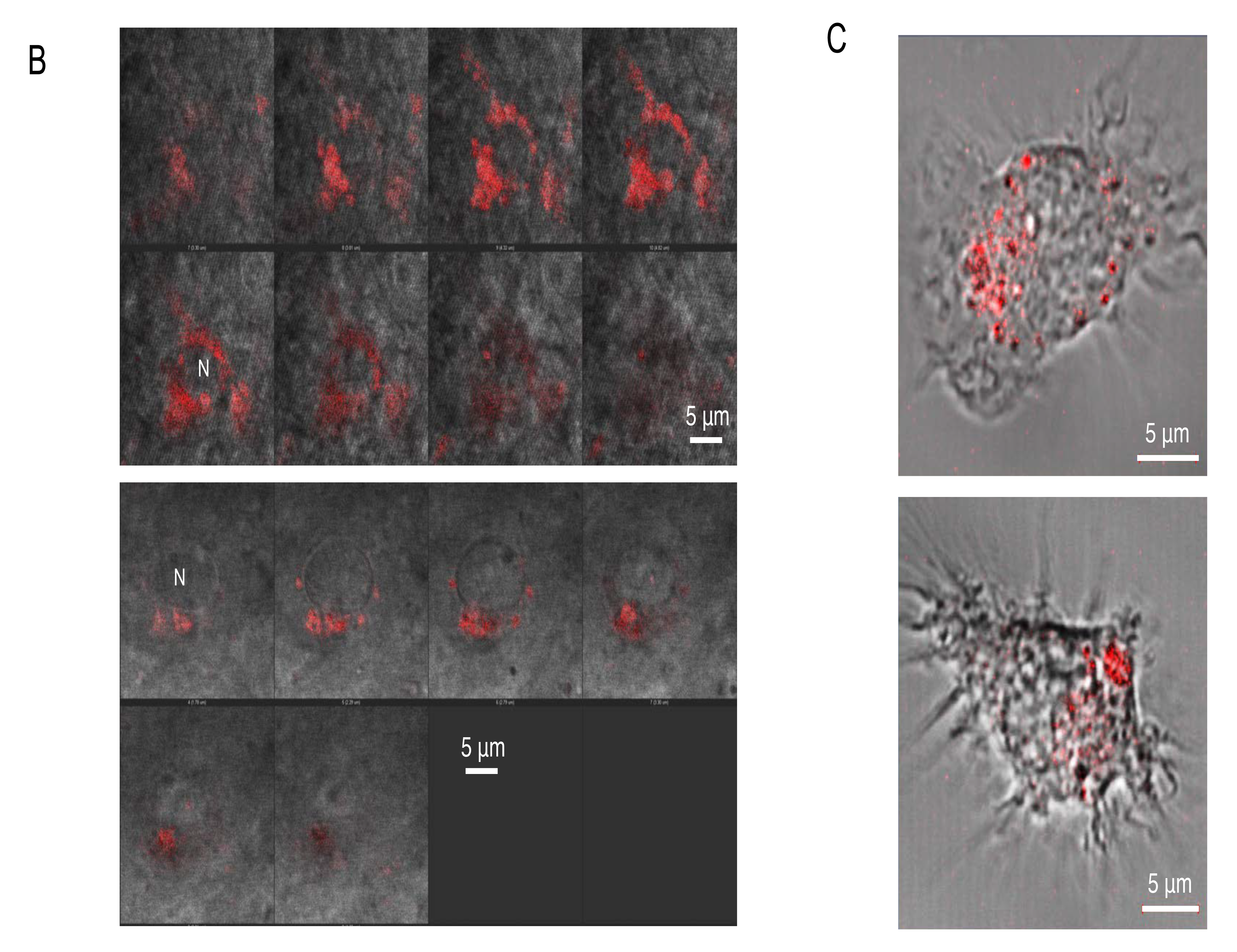
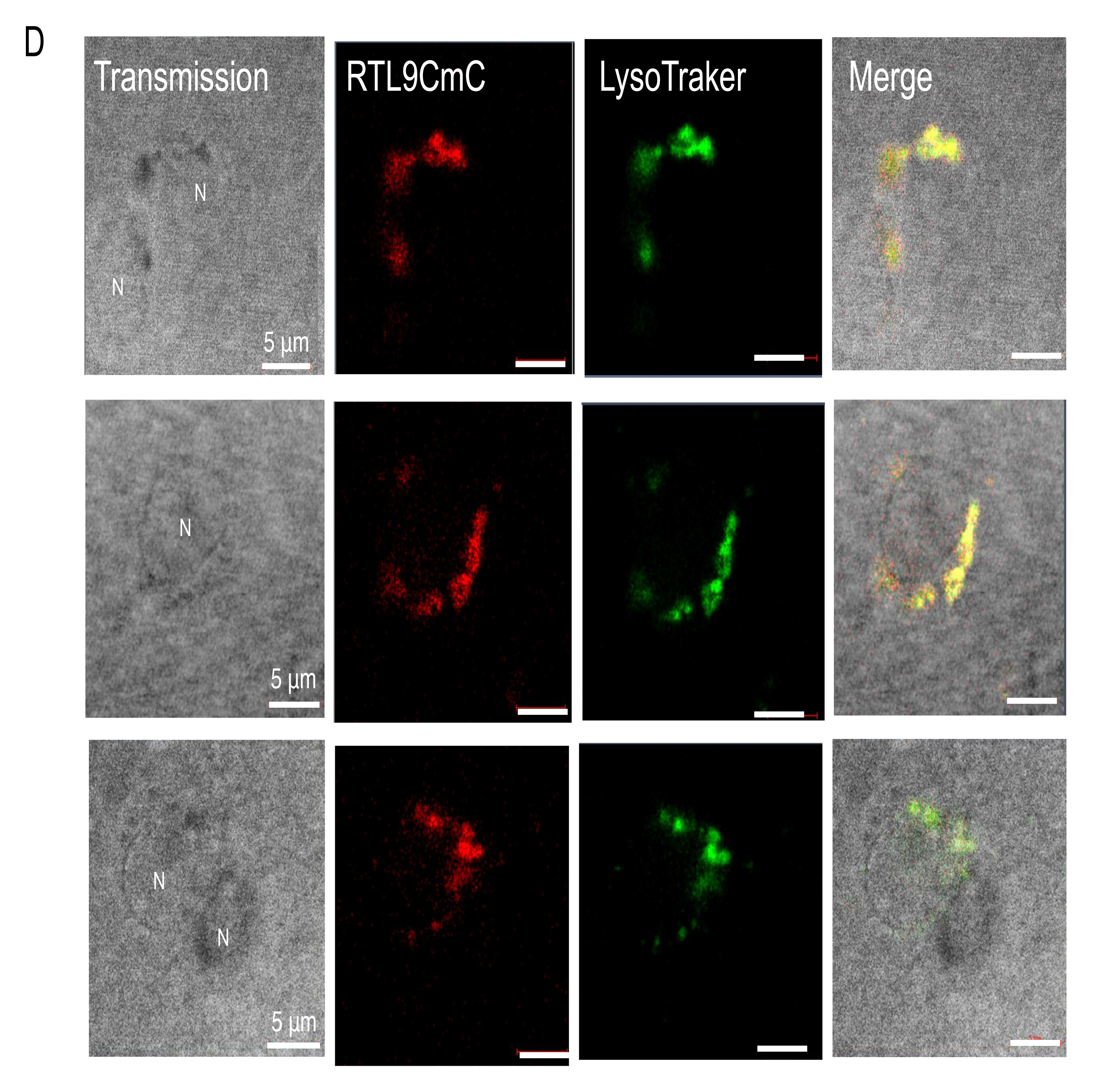
Expression of the RTL9CmC protein in the brain and isolated microglia. **A)** A high expression of RTL9-CmC was detected in the cerebral aqueduct (AQ), commissure of the superior and inferior colliculus (CSC and CIC), periaqueductal gray (PAG), posterior commissure (PC) and superior colliculus (SC), but not in the inferior colliculus (IC), hypothalamus (Ht) or thalamus (Th). The mCherry signals were almost negligible in the WT control. Left: *Rtl9CmC* KI brain (P15). Right: WT control brain (P18). (n>8) **B)** Two sequences of photographs at a 0.5 μm interval indicated that the RTL9CmC protein accumulated in small granules around nuclei in the round cells in the brain (P5). (n=5) **C)** The RTL9CmC protein was detected in both small and large granules in isolated microglia cultured from the P0 brain. (n=2). **D)** The RTL9CmC protein was colocalized with LysoTracker, a specific lysosome marker, in the P9 brain (n=3).

On higher magnification, the signals were detected in a number of small granules (approximately 1 μm diameter or less) that accumulated near the nucleus in round cells (Fig. 2B). We confirmed that these RTL9 expressing cells are microglia because the RTL9CmC protein was detected in similar granules in the isolated microglia cells cultured from P0 neonates (Fig. 2C and Fig. S10) (Lian et al., 2013). In addition, we identified the RTL9 expressing lysosome granules using the lysosome-specific marker, LysoTracker Red NDN99 (Fig. 2D). Thus, it turned out that the RTL9CmC protein localizes in the microglial lysosomes in the brain.

### Involvement of RTL9 in brain innate immunity against fungi

Upon fungal zymosan injection to the brain, RTL9 co-localizes with the incorporated zymosan in microglial lysosomes and promotes its degradation. We reasoned that RTL9 must be functional in brain innate immunity like RTL5 and RTL6 because it also exhibits microglial expression. The RTL5 and RTL6 proteins in the secretory granules of microglia are secreted into the extracellular space and there await their targets, such as dsRNA and LPS, while RTL9 is specific to microglial lysosomes and does not exit into the extracellular space, implying that RTL9 reacts to pathogens that are removed by different mechanisms. We analyzed the response of RTL9 to fungal zymosan because previous reports have shown that zymosan is promptly incorporated into lysosomes (Underhill, 2003). Zymosan is initially recognized by cell surface receptors, such as the TLR2/TLR6 heterodimer and/or other components, then trapped in phagosomes in an LC3-dependent manner and finally degraded in phagolysosomes by fusion with lysosomes (Sanjuan et al. 2007; Martinez et al., 2015; Fiebich et al., 2018; Feldman et al., 2019).

Zymosan is a complex biomolecule containing protein-carbohydrate complex elements such as glucan, mannan, chitin and protein in addition to glycolipid, and therefore exhibits specific autofluorescence patterns (shown in artificial green in Fig. 3) that can be distinguished from other autofluorescence patterns in the brain using the LSM880 Automatic Component Extraction (ACE) function (Fig. S11), as described in our previous report (Irie et al., 2022). Large aggregates of zymosan are also evident in the transmission image (arrows in Fig. S12), so the reliability of its autofluorescence pattern may be verified.

**Figure 3.**
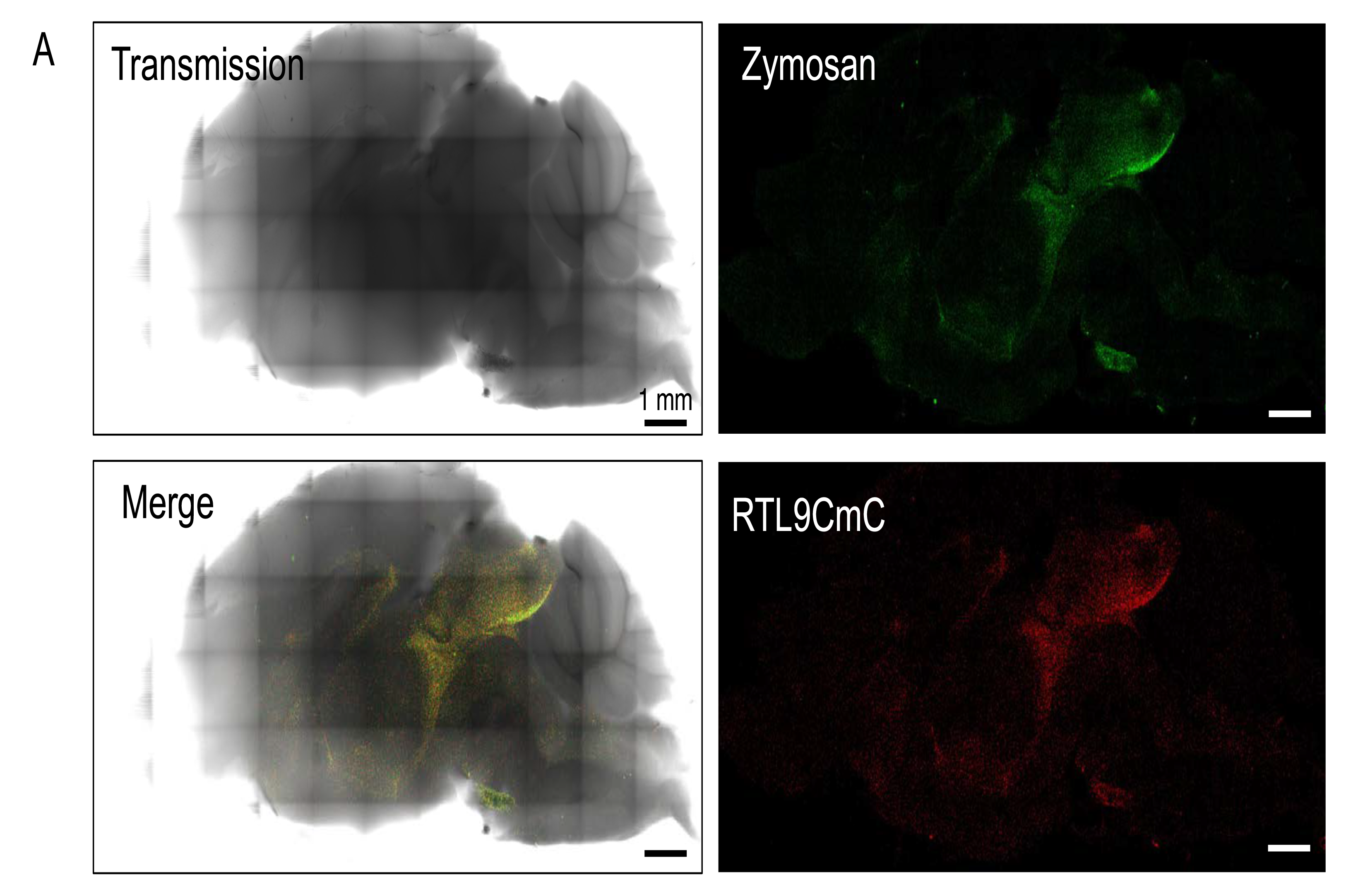
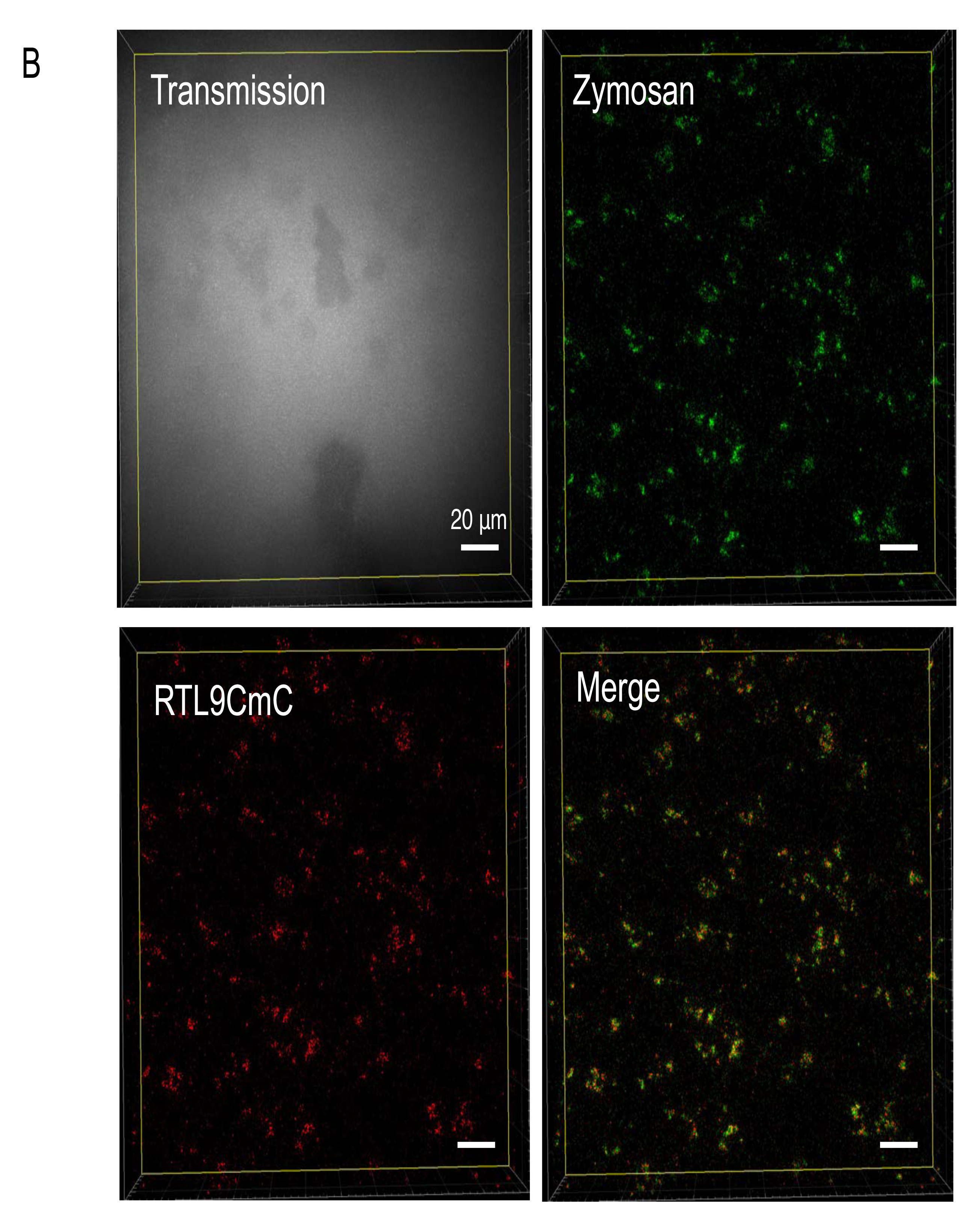
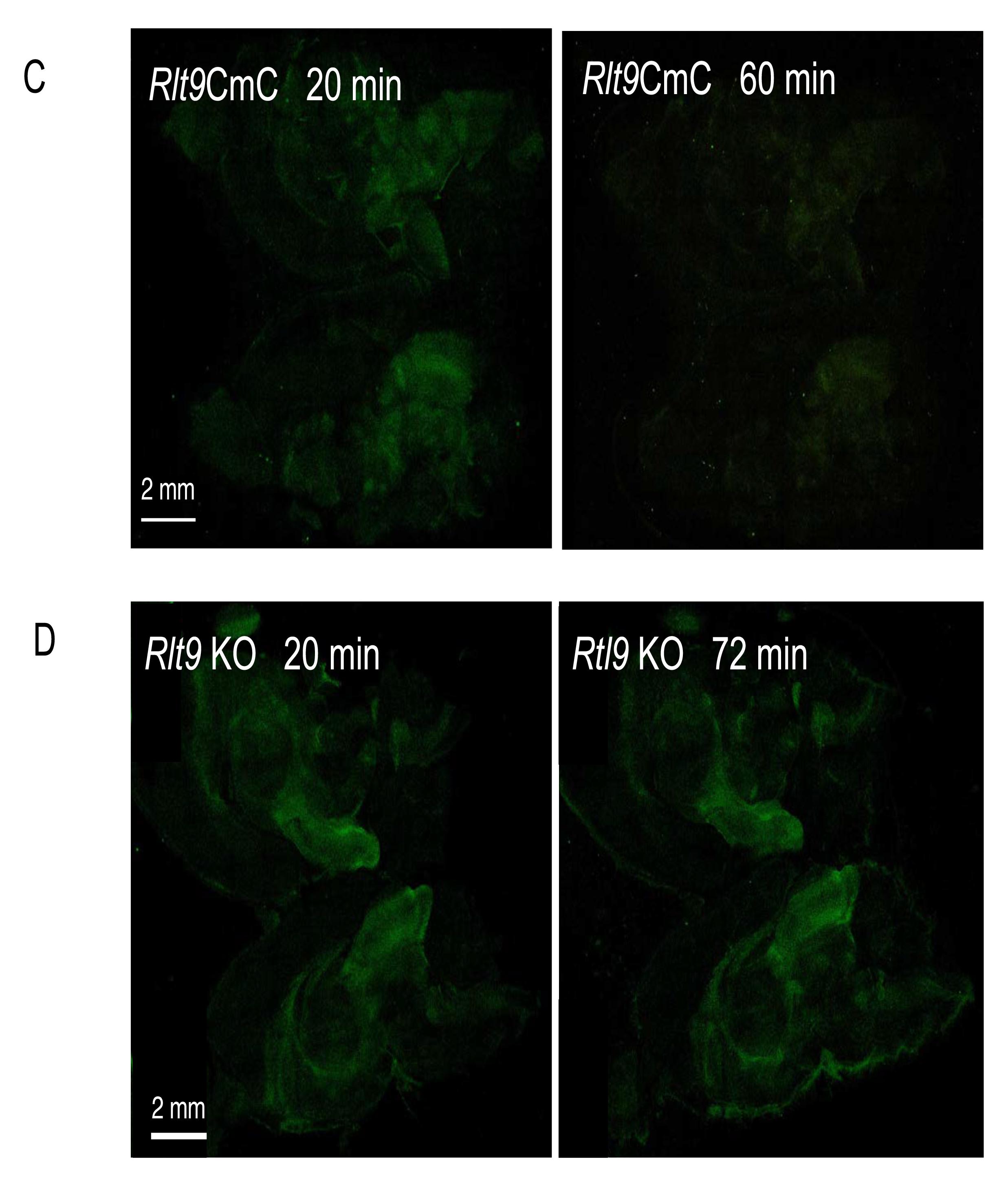
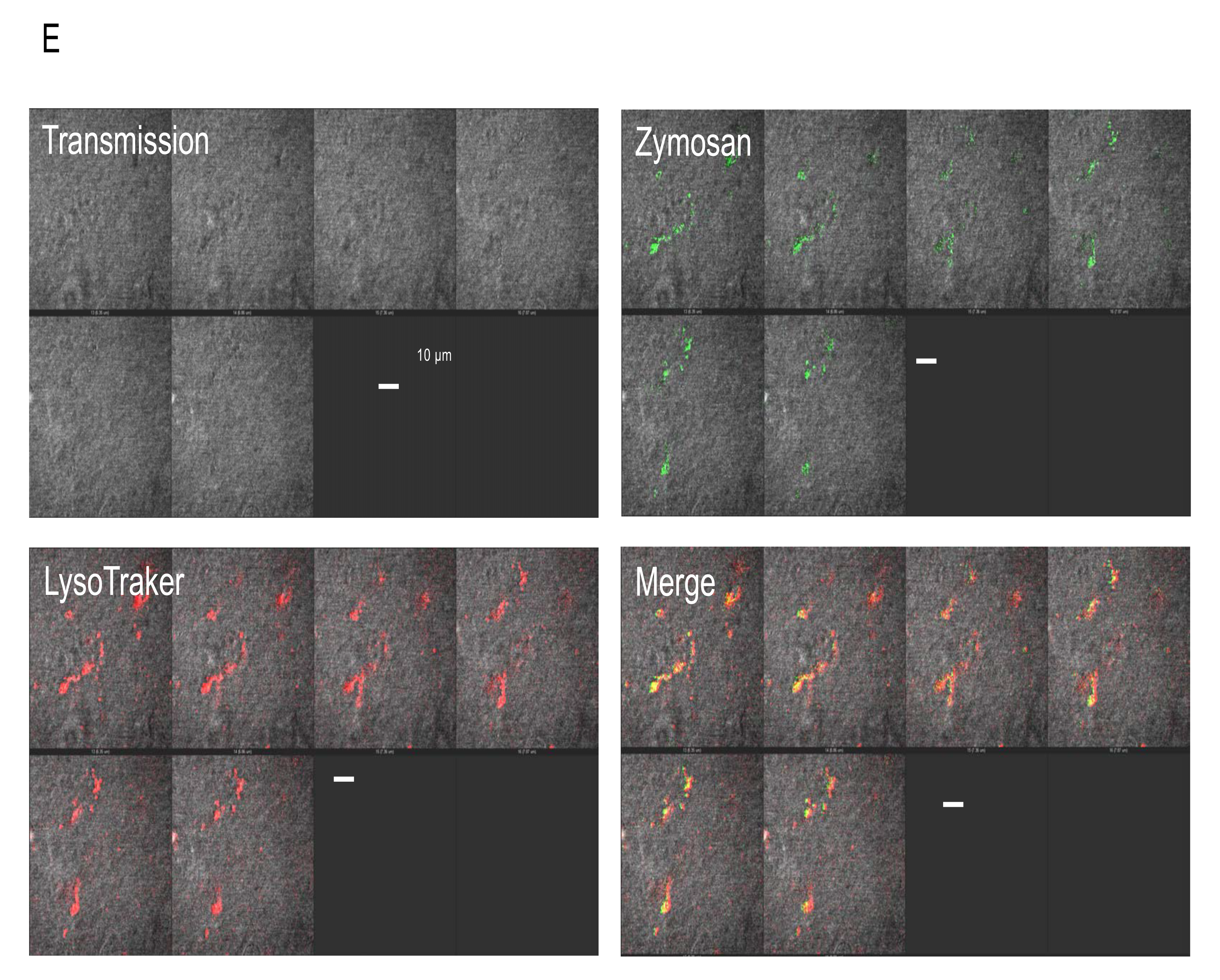
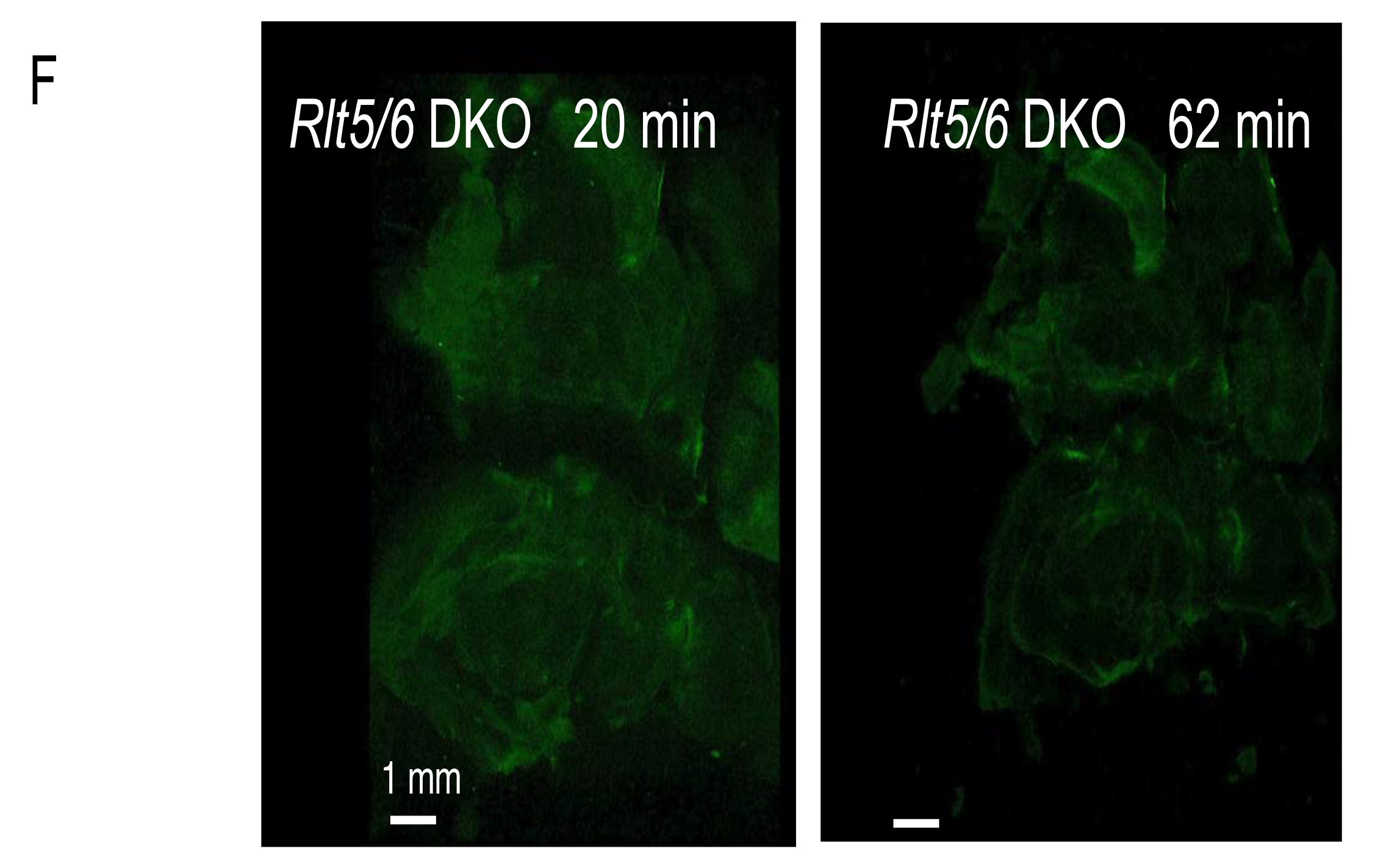
Reaction between the RTL9CmC protein and zymosan. **A)** Colocalization of RTL9CmC (red) and zymosan (green) in the brain. n=5. **B)** A higher magnification image of the midbrain region. n=3. **C)** and **D)** Time course of zymosan degradation in *Rtl9CmC* KI (**C**, 20 and 60 min) and *Rtl9* KO brain (**D**, 20 and 72 min), respectively. n=2. **E)** Lysosomal localization of injected zymosan in *Rtl9* KO brain. 90 min after injection, Lysotracker was added and reacted for 30 min at RT, then a fluorescent image was obtained. n=3. **F)** Normal zymosan degrading activity in *Rtl5/Rtl6* DKO brain. n=3.

Twenty-30 min after the zymosan injection into the brain, we confirmed that zymosan had colocalized with RTL9CmC (Figs. 3A and B). The signal from zymosan was greatly reduced 60 min after the injection, indicating that zymosan had been degraded in the lysosomes (Fig. 3C). We then examined the reaction in the *Rtl9* KO brain and found that the signal intensity remained unchanged even after 72 min (Fig. 3D). This clearly demonstrates that RTL9 is essential for the zymosan degradation reaction in the microglial lysosomes, although it remains unknown how RTL9 is involved and/or what the precise role of RTL9 is in this reaction. We also found that zymosan was normally incorporated into the lysosomes even in the *Rtl9* KO brain (Fig. 3E), suggesting that RTL9 may not be necessary for the fusion between the zymosan-containing phagosomes and lysosomes. We also confirmed that zymosan was normally degraded in the *Rtl5* and *Rtl6* DKO brain (Irie et al., 2022), indicating that the zymosan degradation activity is specific to *Rtl9* (Fig. 3F).

## Discussion

Fungal infections are typically rare yet uniquely dangerous because they are difficult to diagnose and treat. Fungal diseases cause significant morbidity and mortality, particularly in immunocompromised individuals (Brown, 2011; Feldman et al., 2019). This is particularly the case when the fungus invades the brain and central nervous system, such as fungal meningitis. Many fungi in our environment can cause meningitis, including *Aspergillus*, *Candida*, *Cryptococcus*. *Coccidioides* and *Histoplasma*. Recent studies on innate and adaptive immunity have reported that phagocytic cells are essential players in protecting against fungal diseases and that defects in these cells compromise the host’s ability to resist fungal infection (Brown, 2011; Feldman et al., 2019).

In this investigation we have demonstrated that retrovirus-derived *Rtl9* is another microglial gene playing an important role in anti-fungal protection via its ability to degrade zymosan in lysosomes. The *Rtl9* mRNA level is quite low and requires 32 cycles of PCR for its detection even in the brain (Fig. 1B), therefore, the KI mice expressing the RTL9-CmCherry fusion protein from an endogenous locus (Fig. 1C and Fig. S8) were essential for the detection of the RTL9 protein *in vivo*, determination of its location in the microglial lysosomes, and its involvement in the zymosan reaction (Figs. 2 and 3A, B). Finally, in *Rtl9* KO mice it was demonstrated that RTL9 is essential for zymosan degradation in the microglial lysosomes (Figs. 3C and D). We also confirmed that the zymosan degradation activity is specific to RTL9 and not RTL5 or RTL6, other microglial proteins for removal of dsRNA and LPS, respectively (Irie et al., 2022) (Fig. 3F). It is likely that this is the reason why *RTL9* is so robustly well conserved across all eutherian species (dN/dS < 1, Table 1), because protection against fungal infection in the brain provides an evident evolutionary advantage to go along with the protection afforded against viral and bacterial infection by *RTL5* and *RTL6*, respectively (Irie et al., 2022). Considering that *RTL9* is in charge of the eutherian-specific zymosan degradation mechanism, it may well be that *RTL9* is involved in the defense against fungal diseases in the brain.

In contrast to the widely conserved TLR system in the animal kingdom that regulates cytokine and interferon production against various pathogens in a pathogen-dependent manner via each differentiated TLR protein (Hanisch and Kettenmann, 2007; Norris and Kipnis, 2016), RTL9 as wells as RTL5 and RTL6 are apparently novel constituents in the brain innate immune system in eutherians (Irie et al., 2022). Together with the previous study, this work demonstrates that at least three RTL genes are exapted (domesticated) in the current eutherian brain innate immune system and play important roles in the prompt removal of bacteria, virus and fungi in a pathogen-dependent manner. However, the removal of such pathogens is a common activity of microglia/macrophages and not specific to eutherians, suggesting the existence of alternative and/or redundant pathways in non-eutherian organisms and eutherians as well. Therefore, it is also of interest that other retrovirus-derived genes have also been exapted in non-eutherian organisms.

What is the relationship between the TLR system and RTL genes? In the case of zymosan, it is known that the TLR2/TLR6 heterodimer first recognizes zymosan on the cytoplasmic membrane, incorporates it into phagosomes associated with CD14 and then, together with alternative pathways, sends the signal downstream for gene regulation in the course of these processes (Underhill et al., 1999; Underhill, 2003). As RTL9 apparently reacts to zymosan subsequent to fusion of the zymosan-containing phagocytes and lysosomes where RTL9 is localized, it is reasonable to think that the TLR2/TLR6 system is able to function independently of RTL9 and regulate inflammation via cytokine and interferon induction on its own. It was recently suggested that the TLR2/TLR5 heterodimer also induces inflammation when stimulated with zymosan, although TLR5 is known to primarily react to bacterial flagellin (Reuter et al., 2021). Our preliminary experiments confirmed upregulation of *Tnfa*, *Il6* and *Il8* mRNA (Friedland et al., 2001, Bernath et al., 2023) in the *Rtl9* KO microglia in the same manner as control WT microglia, supporting this idea (Fig. S13). It is possible that *Rtl9* itself exhibits upregulation upon zymosan administration, due to the presence of a feedback mechanism. Therefore, more detailed analysis is required to properly elucidate the relationship between the TLR system and *RTL9* as well as the *RTL9* self-regulation mechanism.

What is the role of RTL9 in the zymosan degrading activity, does it act as an enzyme or a modulator? Lysosomes have a variety of hydrolytic enzymes including more than 50 glucosidases (https://www.rndsystems.com/research-area/lysosomal-enzymes). Is RTL9 another lysosomal enzyme? Unfortunately, at present it is not possible to determine whether it has such enzymatic activity or not due to the lack of adequate information.

The retrotransposon/retroviral GAG protein is comprised of three parts, the matrix, capsid and nucleocapsid, and each part is generated by the POL protease. They are structural proteins for virus formation, and therefore have no intrinsic enzymatic activities. Recently, certain GAG-derived genes, such as *ARC*, *PEG10* and *RTL1*, as well as other RTL and Paraneoplastic Ma antigen (PNMA) genes, have attracted attention because they form virus-like particles that are able to mediate cell-cell communication by sending mRNA/siRNAs, and *PEG10* in particular acts as a delivery vehicle for specific mRNAs (Segel et al., 2021). RTL5 and RTL6 are secreted proteins that trap dsRNA and LPS, respectively, in the brain (Irie et al., 2022), implying that they play a role as structural proteins. Therefore, we reason that the GAG-like region of RTL9 forms a complex with zymosan to promote (or modulate) degradation of this complex by an oxidative mechanism. This is because that zymosan is easily solubilized to a high molecular weight polysaccharide via the activity of the hypochlorous acid generated by the heme protein myeloperoxidase (MPO), so it is suggesting that oxidative degradation is the main metabolic pathway of particulate (zymosan) β-glucans (Miura et al., 1996).

BLLF1 comprises 907 aa and is a type 1 membrane protein also known as glycoprotein 350 (gp350). The large N-terminal segment (1-860 aa) extends outside the viral membrane, followed by the transmembrane/cytoplasmic tail regions (861-907 aa) (Kanekiyo et al., Cell 2015). It is known that its extreme N-terminus portion (1-470 aa, domains I, II and III) can bind to CR2, the receptor of Epstein-Barr virus, but it is not essential for this binding reaction because of the presence of alternative proteins that fulfill the same function (Janz et al., 2000). It is structurally well-defined (Szakonyi et al. 2006), but the RTL9 homologous portion (473-780 aa) just after it is functionally and structurally undetermined (Kanekiyo et al. Cell 2015) (Fig. S3).

UL36 consists of 3,164 aa and possesses two leucine-zipper motifs, a proline-glutamine (PQ) repeat, and several ATP-binding sites (McNabb and Courtney, 1992). Its RTL9 homologous portion (787-1141 aa) corresponds to the PQ repeat region, but there are no apparent PQ repeats in RTL9 (Fig. S4). Although there are several PQ motifs throughout the entire RTL9 protein, no PQ motifs are conserved in 10 of the eutherian species (6 PQ motifs are conserved in at least 8 species) (Fig. S1, green dashed boxes). It is reported that the PQ motif on a loop in the cystinosin protein, the lysosomal cystine transporter, is critical for cystinosin transport to lysosomes (Cherqui et al., 2001; Ponting et al. 2001). It is therefore possible that certain PQ motif(s) play a role in RTL9 transport to lysosomes. Unfortunately, there is no RTL9 3D structure prediction afforded by Alphafold 2 except the GAG-like region. Therefore, further studies are obviously required to elucidate the biochemical role of RTL9 in the zymosan degradation reaction in detail.

In this study the importance of the RTL9 C-terminal part in zymosan degradation has been demonstrated. This activity potentially arises out of the entire sushi-ichi GAG (the maximal GAG-like region) or possibly comprises of the latter half of UL36-like region and the minimal GAG-like region (Fig. 1A and Figs. S1 and S4 to S6). However, it is also possible that the remaining large regions contribute to the enzymatic zymosan degrading activity in combination with the C-terminal region. We confirmed that the incorporated zymosan ultimately reached the microglial lysosomes even in *Rtl9* KO mice, meaning that the C-terminal portion is not essential for either zymosan incorporation into phagosomes or the phagolysosome fusion process (Fig. 3D). However, it is also possible that the other regions may be required for these processes.

Zymosan degradation is unique because it is carried out via LC3-mediated phagocytosis in association with several autophagy components (Sanjuan et al. 2007; Martinez et al., 2015; Fiebich et al., 2018, Feldman et al., 2019). Further studies are required to elucidate the entire function(s) of the RTL9 protein during these processes including both fusion of the zymosan-incorporated phagosomes with lysosomes and zymosan degradation by generating another KO mice in which the entire RTL9 sequence is deleted.

Among the 11 RTL genes, *PEG10*, *RTL1* (aka *PEG11*) and *LDOC1* (aka *SIRH7* and *RTL7*) play essential but different roles in the placenta (Ono et al., 2006; Sekita et al., 2008; Kagami et al., 2008; Naruse et al., 2014; Kitazawa et al., 2017; Shiura et al., 2021), while *RTL5*, *RTL6* (Irie et al., 2022) and *RTL9* (this study) play important but different roles in microglia. Microglia originate in early development from the yolk sac and not from the bone marrow as do most macrophages (Ginhoux et al., 2010 and 2013). Therefore, *RTL9* provides considerable support for a central hypothesis in our work, which is that extraembryonic tissues, such as the yolk sac and placenta, have served as incubation sites for the birth of newly acquired genes from retrotransposons/retroviruses in the course of eutherian evolution (Kaneko-Ishino and Ishino, 2015; Irie et al., 2022).

## Materials and Methods

### Mice

All of the animal experiments were reviewed and approved by the Institutional Animal Care and Use Committee of Tokai University and Tokyo Medical and Dental University (TMDU) and were performed in accordance with the Guideline for the Care and Use of Laboratory Animals of Tokai University and TMDU.

### Comparative genome analysis

The sushi-ichi GAG (AAC33525.1) and mouse RTL9 (NP_001035524.2) protein sequences were obtained from NCBI, while the amino acid identity and similarity were calculated using the EMBOSS Water program (https://www.ebi.ac.uk/Tools/psa/emboss_water/) and EMBOSS needle program (https://www.ebi.ac.uk/Tools/psa/emboss_needle/) in the default mode. The orthologues of RTL9 were identified by a search of the NCBI Gene database (https://www.ncbi.nlm.nih.gov/gene/) using RTL9 (and RGAG1) as the query terms. The RTL9 coding sequences used for the pairwise dN/dS analysis (Table 1) and the amino acid alignment (Fig. S1) were as follows: Mouse (*Mus musculus*): NM_001040434.2; Rat (*Rattus norvegicus*): XM_003752123.5; Human (*Homo sapiens*): NM_001385449.1; Chimpanzee (*Pan troglodytes*): XM_016942939.3; Dog (*Canis lupus familiaris*): XM_038449037.1; Horse (*Equus caballus*): XM_005614417.3; Elephant (*Loxodonta Africana*): XM_003414859.3; Manatee (*Trichechus manatus latirostris*): XM_023734430.1; Armadillo (*Dasypus novemcinctus*): XM_023582615.1; Sloth (*Choloepus didactylus*): XM_037822012.1. The RTL9 coding sequenses were translated to protein sequences using the EMBOSS Transeq (https://www.ebi.ac.uk/Tools/st/emboss_transeq/) program. The amino acid alignment in the ten eutherian species was constructed using the Clustal Omega program (https://www.ebi.ac.uk/Tools/msa/clustalo/) in the default mode. The conserved domains in RTL9 were identified by means of the NCBI Conserved Domains Database (CDD) (https://www.ncbi.nlm.nih.gov/Structure/cdd/wrpsb.cgi).

### Estimation of the pairwise dN/dS ratio

The nonsynonymous/synonymous substitution rate ratio (dN/dS) was estimated with CodeML (runmode: −2) in PAML (Xu and Yang, 2013). An amino acid sequence phylogenic tree was constructed with MEGA7 (Kumar et al., 2016) using the Maximum Likelihood method with the JTT matrix-based model. The codon alignment of cDNA was created with the PAL2NAL program (www.bork.embl.de/pal2nal/) (Suyama et al., 2006). The RTL9 coding sequences used for the pairwise dN/dS ratio analysis (Table 1) are described above.

### RT-PCR

Total RNA samples were prepared from adult tissues (11w) using ISOGEN (Nippon Gene). The cDNA was synthesized from 1 μg of total RNA using Superscript III reverse transcriptase (Invitrogen) with an oligo-dT primer. For RT-PCR, 10 ng cDNA in a 25 μl reaction mixture containing 1 X ExTaq buffer (TaKaRa), 200 μM each of NTP and 800 nM of primers along with 0.625 units *ExTaq* HS (TaKaRa) were subjected to 32 cycles at 96°C for 15 s, 65°C for 30s and 72°C for 30s in a GeneAmp PCR System 2400 (Perkin-Elmer). The primer sequences used were as follows: *β*-actin-forward 5’- AAGTGTGACGTTGACATCC-3’ and *β*-actin-reverse 5’- GATCCAACTCTGCTGGAAGG-3’; *Rtl9*-forward 5’-TCACCTACATGCCTGTGACC- 3’ and *Rtl9*-reverse 5’-CAACAACACCACATTGTTACGG-3’.

### Generation of the *Rtl9-mCherry* knock-in mice

*Rtl9*-*mCherry* knock-in mice were generated by pronuclear microinjection of the CRISPR/Cas system (see Fig. 1C), essentially as described in a previous report (Aida et al., 2015), using a plasmid targeting vector. The crRNA was designed and synthesized with the target sequence: 5’-TACAAACAAGTAGTACTCCT-3’ (Fasmac). The plasmid for mCherry insertion was constructed with 1.5 kb long 5’ and 3’ arms amplified from the C57BL/6N genome using PrimeSTAR GXL DNA Polymerase (TaKaRa). The 5’ arm is the genomic sequence upstream of the stop codon of Rtl9 and the 3’ arm is downstream of the predictive Cas9 cut site. The C-terminus of Rtl9 was fused to a 4x GGS linker (ggaggatcaggaggatcaggaggatcaggaggatca)-attached mCherry by means of a cloning enzyme (In-Fusion® HD Cloning Kit, Takara Bio). The purified targeting vector was assessed for its quality by Sanger sequencing and injected into mouse pronuclei at the final concentration of 10 ng/μl. Just before injection, the targeting vector was mixed with other CRISPR/Cas system components, including the Cas9 protein (a final concentration of 30 ng/μl), crRNA (8.7 ng/μl), tracrRNA (14.3 ng/μl) and injected into the pronuclei of mouse zygotes produced by *in* vitro fertilization using C57BL/6N mice. Zygotes that survived the microinjection procedure were cultured in KSOM at 37°C under 5% CO2 in air and transferred into the ampulla of the oviduct of pseudopregnant ICR females. Genomic modification of the mutant mice was confirmed using 3 sets of PCR primers (5’- GGAATGATGTCCACGCCACTA-3’ and 5’- CTTCAGCTTCAGCCTCTGCT-3’, 5’-CCTGTCCCCTCAGTTCATGT-3’ and 5’- CCTAGACTATTGGACCAGAGG-3’, and 5’-GGAATGATGTCCACGCCACTA-3’ and 5’-CCTAGACTATTGGACCAGAGG-3’).

### Generation of the *Rtl9* knock-out mice

*Rtl9* knock-out mice **we**re generated using a pair of crRNAs and a single-stranded oligo donor DNA (ssODN). To cut off the C-terminal genomic region of Rtl9, a pair of crRNAs targeting 5’-TCATAGCTGCAAATTGTGCA-3’ and 5’- AGAAATCACATAAGATTCCA-3’ was synthesized (Fasmac). The 162-base ssODN was composed of a 3x stop codon (3x stop) and 5’ and 3’ homology arms having 75 bases each (Hokkaido System Science). Just before injection, CRISPR/Cas system components including the Cas9 protein (a final concentration of 30 ng/μl), a pair of crRNAs (8.7 ng/μl), tracrRNA (14.3 ng/μl) and ssODN (15 ng/μl) were mixed and injected into the pronuclei of mouse zygotes produced by *in vitro* fertilization using C57BL/6N mice. The zygotes that survived the microinjection procedure were cultured in KSOM at 37°C under 5% CO2 in air and transferred into the ampulla of the oviduct of pseudopregnant ICR females. Genomic modification of the mutant mice was confirmed using a set of PCR primers (5’- GAGAACACCAGCTTCTAGAGC-3’ and 5’- GGGAGTTCAGAACCTCATACAC-3’).

### Imaging using Confocal Laser Scanning Fluorescence Microscopy

Fresh brain hemispheres from *Rtl9*-*mCherry* KI mice were used for analysis with a ZEISS LSM880 (ZEISS, Germany) without fixation. Samples were covered with 10% glycerol solution for protection from drying. The samples were observed using a Plan-Apochromat lens (10x, numerical aperture =0.45, M27, ZEISS) and a C-Apochromat lens (63x numerical aperture =1.2 Water, ZEISS). The tiling with lambda-mode images was obtained using the following settings: pixel dwell: 1.54 μs; average: line 4; master gain: ChS: 1250, ChD: 542; pinhole size: 33 µm; filter 500 – 696 nm; beam splitter: MBS 458/514; lasers: 514 nm (Argon 514), 0.90%. For the tiling-scan observations, the tiling images were captured as tiles: 84, overlap in percentage: 10.0, tiling mode: rectangular grid, in size: x; 15442.39 µm, y; 9065.95 µm. Spectral unmixing and processing of the obtained images were conducted using ZEN imaging software (Carl Zeiss Microscopy, Jena Germany). The spectrum from the mCherry protein (Maximum peak emission fluorescence wavelength: 610 nm) detected in *Rtl9-mCherry* KI brain was distinguished from Af610 nm using the peak shape, such as the width and/or co-existence of a second peak. The relative intensity of RTL9-mCherry (red), zymosan autofluorescence (green) and LysoTraker Red NDN99 (red in Fig. 3E or green in Fig. 2D) signals along the x-axis of the brain (from the olfactory bulb to cerebellum region) was calculated from 3D scanning data. The total intensity of each signal on and above each y-axis was summed, divided by the transmission signal and presented as the relative signal intensity on the y-axis in this figure.

### Zymosan injection to the brain

*Rtl9*-*mCherry* KI, *Rtl9* KO and *Rtl5* and *Rtl6* double KO mice (Irie et al., 2022) (P5 to 3w of age) were used for the injection experiments after being anesthetized with isoflurane. Ten µl of 10 ng/ml solution of Zymosan (NOVUS BIOLOGICALS, NPB2- 26233, 1/5000 dilution) were injected using 1ml insulin syringes and a 26 G needle. One min after the injection, the needle was pulled out and kept out for 5 min, then the fresh brain was dissected out in cold PBS solution. The inner surface of the brain hemispheres was analyzed with a ZEISS LSM880 (ZEISS, Germany).

### Lysosome staining with LysoTracker

A 1/50 dilution of LysoTrackerTM Red DND-99 (ThermoFisher Scientific, L7528, 1mM) was used for lysosome staining of fresh brains. A 100 µl solution was placed beneath the brain hemispheres and reacted for 30 min at RT. The LysoTracker signal (emission peak at 590 nm) was separated from various kinds of autofluorescence (Af) in the brain using the LSM880 ACE function.

### Competing interests

The authors declare that the research was conducted in the absence of any commercial or financial relationships that could be construed as a potential conflict of interest.

## Author contributions

F.I., J.I., M.I., A.M., M.N. and T.K.-I. performed the experiments and analyzed the data. A.M., T.S. and Y.H. generated *Rtl9-mCherry* KI and *Rtl9* KO mice. F.I. and T.K.-I. designed the study and wrote the manuscript. All authors agree to be accountable for the content of the work.

## Funding

This work was supported by funding program for Next Generation World-Leading Researchers (NEXT Program LS112) and Grants-in-Aid for Scientific Research (C) (17K07243 and 21K06127) from Japan Society for the Promotion of Science (JSPS) to T.K.-I, Grants-in-Aid for Scientific Research (S) (23221010) and (A) (16H02478 and 19H00978) from JSPS to F.I., Nanken Kyoten Program, Medical Research Institute, Tokyo Medical and Dental University (TMDU) to T.K.-I. and F.I. The funders had no role in study design, data collection and analysis, decision to publish, or preparation of the manuscript.

## Acknowledgements

We thank Takayasu N at Tokai University and Tokai University Support Center for Medical Research and Education for technical assistance of animal breeding. We also thank Usami T for technical assistance in generating *Rtl9-mCherry* KI and *Rtl9* KO mice. Pacific Edit reviewed the manuscript prior to submission.

## Notes

### Competing Interest Statement

The authors have declared no competing interest.

## References

Aida, T., Chiyo, K., Usami, T., Ishikubo, H., Imahashi, R.,Wada, Y., Tanaka, K. F., Sakuma, T., Yamamoto, T. and Tanaka, K. (2015). Cloning-free CRISPR/Cas system facilitates functional cassette knock-in in mice. Genome Biol. 16, 87. doi:10.1186/s13059-015-0653-x

Akira, S., Uematsu S. and Takeuchi O. (2006). Pathogen Recognition and Innate Immunity. Cell 124, 783–801. doi:10.1016/j.cell.2006.02.015

Bernath, A. K., Murray T., E., Yang, S., Gibon, J. and Klegeris, A. (2023). Microglia secrete distinct sets of neurotoxins in a stimulus-dependent manner. Brain Res. 1807, 148315. https://doi.org/10.1016/j.brainres.2023.148315

Brandt, J., Schrauth, S., Veith, A.-M., Froschauer, A., Haneke, T., Schultheis, C., Gessler, M., Leimeister, C. and Volff, J.-N. (2005). Transposable elements as a source of genetic innovation: expression and evolution of a family of retrotransposon-derived neogenes in mammals. Gene 345, 101–111. doi:10.1016/j.gene.2004.11.022

Brown, G. D. (2011). Innate Antifungal Immunity: The Key Role of Phagocytes. Annu. Rev. Immunol. 29, 1–21. doi: 10.1146/annurev-immunol-030409-101229

Cherqui, S., Kalatzis, V., Trugnan, G. and Antignac, C. (2001). The targeting of cystinosin to the lysosomal membrane requires a tyrosine-based signal and a novel sorting motif. J. Biol. Chem. 276, 13314–13321. doi:10.1074/jbc.m010562200

Chou, M.-Y., Hu, M.-C., Chen, R.-Y., Hsu, C.-L., Lin, T.-Y., Tan, M.-J., Lee, C.-Y., Kuo, M.-F., Huang, P.-H, Wu, V.-C. et al. (2022). RTL1/PEG11 imprinted in human and mouse brain mediates anxiety-like and social behaviors and regulates neuronal excitability in the locus coeruleus.

Edwards, C. A., Mungall, A. J., Matthews, L., Ryder, E., Gray, D. J., Pask, A. J., Shaw, G., Graves, J. A., Rogers, J., SAVOIR consortium, Dunham, I., Renfree, M. B., Ferguson-Smith, A. C. (2008). The evolution of the *DLK1-DIO3* imprinted domain in mammals. PLoS Biol 6, e135.

Feldman, M. B., Vyas, J. M. and Mansour M. K. (2019). It takes a village: Phagocytes play a central role in fungal immunity. Semin Cell Dev Biol 89, 16–23. doi: 10.1016/j.semcdb.2018.04.008

Fiebich, B. L., Batista, C. R. A., Saliba, S. W., Yousif, N. M. and de Oliveira, A. C. P. (2018). Role of microglia TLRs in neurodegeneration. Front. Cell. Neurosci. 12, 329. doi:10.3389/fncel.2018.00329

Friedland, J. S., Constantin, D., Shaw, T. C. and Stylianou E. (2001). Regulation of interleukin-8 gene expression after phagocytosis of zymosan by human monocytic cells. J. Leuko. Biol.70, 447–454.

Hanisch, U.-K. and Kettenmann, H. (2007). Microglia: active sensor and versatile effector cells in the normal and pathologic brain. Nat. Neurosci. 10, 1387–1394. doi:10.1038/nn1997

Imakawa, K., Kusama, K., Kaneko-Ishino, T., Nakagawa, S., Kitao, K., Miyazawa, T. and Ishino, F. (2022). Endogenous Retroviruses and Placental Evolution, Development, and Diversity. Cells 11, 2458. doi: 10.3390/cells11152458.

Irie, M., Yoshikawa, M., Ono, R., Iwafune, H., Furuse, T., Yamada, I., Wakana, S., Yamashita, Y., Abe, T., Ishino. F. and Kaneko-Ishino T. (2015). Cognitive function related to the *Sirh11/Zcchc16* gene acquired from an LTR retrotransposon in eutherians. PLoS Genet 11(9):e1005521. doi: 10.1371/journal.pgen.1005521.

Irie, M., Itoh, J., Matsuzawa, A., Ikawa M., Kiyonari, H., Kihara, M., Suzuki T., Hiraoka, Y., Ishino, F. and Kaneko-Ishino, T. (2022). Retrovirus-derived *RTL5* and *RTL6* genes are novel constituents of the innate immune system in the eutherian brain. Development 149, dev200976. doi:10.1242/dev.200976

Ginhoux, F., Greter, M., Leboeuf, M., Nandi, S., See, P., Gokhan, S., Mehler, M. F., Conway, S. J., Ng, L. G., Stanley, E. R. et al. (2010). Fate mapping analysis reveals that adult migroglia derive from primitive macrophages. Science 330, 841–845. doi:10.1126/science.1194637

Ginhoux, F., Lim, S., Hoeffel, G., Low, D. and Huber, T. (2013). Origin and differentiation of microglia. Front. Cell. Neurosci. 7, 45. doi:10.3389/fncel.2013.00045

Ioannides, Y., Lokulo-Sodipe, K., Mackay, D. J. G., Davies, J. H. and Temple, I. K. (2014). Temple syndrome: improving the recognition of an underdiagnosed chromosome 14 imprinting disorder: an analysis of 51 published cases. J. Med. Genet. 51, 495–501. doi:10.1136/jmedgenet-2014-102396

Janz, A., Oezel, M., Kurzeder, C., Mautner, J., Pich, D., Kost, M., Hammerschmidt, W. and Delecluse, H. J. (2000). Infectious Epstein-Barr virus lacking major glycoprotein BLLF1 (gp350/220) demonstrates the existence of additional viral ligands. J. Virol. 74,10142–10152. doi: 10.1128/jvi.74.21.10142-10152.2000.

Kagami, M., Sekita, Y., Nishimura, G., Irie, M., Kato, F., Okada, M., Yamamori, S., Kishimoto, H., Nakayama, M., Tanaka, Y. et al. (2008). Deletions and epimutations affecting the human 14q32.2 imprinted region in individuals with paternal and maternal upd(14)-like phenotypes. Nat. Genet. 40, 237–242. doi:10.1038/ng.2007.56

Kagami, M., Kurosawa, K., Miyazaki, O., Ishino, F., Matsuoka, K. and Ogata, T. (2015). Comprehensive clinical studies in 34 patients with molecularly defined UPD(14)pat and related conditions (Kagami-Ogata syndrome). Eur. J. Hum. Genet. 23, 1488–1498. doi:10.1038/ejhg.2015.13

Kanekiyo, M, Bu, W., Joyce, M. G., Meng, G., Whittle, J. R. R., Baxa, U., Yamamoto, T., Narpala, S., Todd, J-P., Rao, S. S., et al. (2015). Rational Design of an Epstein-Barr Virus Vaccine Targeting the Receptor-Binding Site. Cell 162, 1090–1100. doi:10.1016/j.cell.2015.07.043

Kaneko-Ishino, T. and Ishino, F. (2012). The role of genes domesticated from LTR retrotransposons and retroviruses in mammals. Front. Microbiol. 3, 262. doi:10.3389/fmicb.2012.00262

Kaneko-Ishino, T. and Ishino, F. (2015). Mammalian-specific genomic functions: Newly acquired traits generated by genomic imprinting and LTR retrotransposonderived genes in mammals. Proc. Jpn. Acad. Ser. B Phys. Biol. Sci. 91, 511–538. doi:10.2183/pjab.91.511

Kim, A., Terzian, C., Santamaria, P., Pélisson, A., Purd’homme, N., and Bucheton, A. (1994). Retroviruses in invertebrates: The gypsy retrotransposon is apparently an infectious retrovirus of Drosophila melanogaster. Proc. Natl. Acad. Sci. USA, 91, 1285–1289.

Kitazawa, M., Tamura, M., Kaneko-Ishino, T., and Ishino, F. (2017). Severe damage to the placental fetal capillary network causes mid to late fetal lethality and reduction of placental size in *Peg11/Rtl1* KO mice. Genes Cells, 22, 174–188. doi: 10.1111/gtc.12465.

Kitazawa, M., Hayashi, S., Imamura, M., Takeda, S., Oishi, Y., Kaneko-Ishino, T. and Ishino, F. (2020). Deficiency and overexpression of Rtl1 in the mouse cause distinct muscle abnormalities related to Temple and Kagami-Ogata syndromes. Development 147, dev185918. doi:10.1242/dev.185918

Kitazawa, M., Sutani, A., Kaneko-Ishino, T. and Ishino, F. (2021). The role of eutherian-specific RTL1 in the nervous system and its implications for the Kagami-Ogata and Temple syndromes. Genes Cells 26, 165–179. doi:10.1111/gtc.12830

Kumar, S., Stecher, G. and Tamura, K. (2016). MEGA7: Molecular Evolutionary Genetics Analysis Version 7.0 for Bigger Datasets. Mol. Biol. Evol. 33, 1870–1874. doi:10.1093/molbev/msw054

Lian, H., Roy, E. and Zheng, H. (2016). Protocol for primary microglial culture preparation. Bio. Protoc. 6, e1989. doi:10.21769/BioProtoc.1989

Lim, E. T., Raychaudhuri, S., Sanders, S. J., Stevens, C., Sabo, A., MacArthur, D. G., Neale, B. M., Kirby, A., Ruderfer, D. M., Fromer, M., et al. (2013). Rare complete knockouts in humans: population distribution and significant role in autism spectrum disorders. Neuron 77, 235–242. doi:10.1016/j.neuron.2012.12.029

Luzio, J. P., and Pryor, P. R. and Bright, N. A. (2007). Lysosomes: fusion and function. Nat. Rev. Mol. Cell Biol. 8, 623–632. doi:10.1038/nrm2217

Martinez, J., Malireddi, R. K., Lu, Q., Cunha, L. D., Pelletier, S., Gingras, S., Orchard, R., Guan, J.-L., Tan, H., Peng, J., Kanneganti. T.-D., Virgin, H. W. and Green D. R. (2015). Molecular characterization of LC3-associated phagocytosis reveals distinct roles for Rubicon, NOX2 and autophagy proteins. Nat. Cell. Biol. 17, 893–906. doi:10.1038/ncb3192

McNabb, D. S. and Courtney, R. J. (1992). Analysis of the UL 36 Open Reading Fame Encoding the Large Tegument Protein (ICP1/2) of Herpes Simplex Virus Type I. J. Virol. 66, 7581–7584.

Miura, N., N., Ohno, N., Adachi, Y. and Yadome, T. (1996). Characterization of Sodium Hypochlorite Degradation of β-Glucan in Relation to Its Metabolism *in Vivo*. Chem. Pharm. Bull. 44, 2137–2141. doi:10.1248/cpb.44.2137

Naruse, M., Ono, R., Irie, M., Nakamura, K., Furuse, T., Hino, T., Oda, K., Kashimura, M., Yamada, I., Wakana, S. et al. (2014). Sirh7/Ldoc1 knockout mice exhibit placental P4 overproduction and delayed parturition. Development 141, 4763–4771. doi:10.1242/dev.114520

Norris, G. T. and Kipnis, J. (2018). Immune cells and CNS physiology: Microglia and beyond. J. Exp. Med. 216, 60–70. doi:10.1084/jem.20180199

Ono, R., Kobayashi, S., Wagatsuma, H., Aisaka, K., Kohda, T., Kaneko-Ishino, T. and Ishino, F. (2001). A retrotransposon-derived gene, PEG10, is a novel imprinted gene located on human chromosome 7q21. Genomics 73, 232–237. doi: 10.1006/geno.2001.6494.

Ono, R., Nakamura, K., Inoue, K., Naruse, M., Usami, T., Wakisaka-Saito, N., Hino, T., Suzuki-Migishima, R., Ogonuki, N., Miki, H. et al. (2006). Deletion of Peg10, an imprinted gene acquired from a retrotransposon, causes early embryonic lethality. Nat. Genet. 38, 101–106. doi:10.1038/ng1699

Ponting, C. P., Mott, R., Bork, P. and Copley, R. R. (2001). Novel protein domains and repeats in Drosophila melanogaster: insights into structure, function, and evolution. Genome Res. 11, 1996–2008. doi:10.1101/gr.198701

Reuter, S., Herold, K., Domrös, J. and Mrowka, R. (2021). Toll-like receptor 5 as a novel receptor for fungal zymosan. bioRxiv. doi: https://doi.org/10.1101/2021.12.23.473960

Salazar, F. and Brown, G, D, (2018). Antifungal Innate Immunity: A Perspective from the Last 10 Years. J Innate Immun 10, 373–397. doi: 10.1159/000488539

Sanjuan, M. A., Dillon, C. P., Tait, S. W. G., Moshiach, S., Dorsey, F., Connell, S., Komatsu, M., Tanaka, K., Cleveland, J. L., Withoff, S. and Green, D. R. (2007). Toll-like receptor signalling in macrophages links the autophagy pathway to phagocytosis. Nature 450, 1253–1257. doi:10.1038/nature06421

Segel, M., Lash, B., Song, J., Ladha, A., Liu, C. C., Jin, X., Mekhedov, S. L., Macrae, R. K., Koonin, E. V. and Zhang, F. (2021). Mammalian retrovirus-like protein PEG10 packages its own mRNA and can be pseudotyped for mRNA delivery. Science 373, 882–889. doi: 10.1126/science.abg6155

Sekita, Y., Wagatsuma, H., Nakamura, K., Ono, R., Kagami, M., Wakisaka, N., Hino, T., Suzuki-Migishima, R., Kohda, T., Ogura, A. et al. (2008). Role of retrotransposon-derived imprinted gene, Rtl1, in the feto-maternal interface of mouse placenta. Nat. Genet. 40, 243–248. doi:10.1038/ng.2007.51

Shiura, H., Ono, R., Tachibana, S., Kohda, T., Kaneko-Ishino, T. and Ishino F. (2021). PEG10 viral aspartic protease domain is essential for the maintenance of fetal capillary structure in the mouse placenta. Development 148, dev199564. doi:10.1242/dev.199564

Song, S.U., Gerasimova, T., Kurkulos, M., Boeke, J.D. and Corces, V.G. (1994). An env-like protein encoded by a Drosophila retroelement: Evidence that gypsy is an infectious retrovirus. Genes Dev. 8, 2046–2057.

Suyama, M., Torrents, D. and Bork, P. (2006). PAL2NAL: robust conversion of protein sequence alignments into the corresponding codon alignments. Nucleic Acids Res. 34, W609–W612. doi:10.1093/nar/gkl315

Suzuki, S., Ono, R., Narita, T., Pask, A. J., Shaw, G., Wang, C., Kohda, T., Alsop, A. E., Graves M. J. A., Kohara, Y., Ishino, F., Renfree, M. B. and Kaneko-Ishino, T. (2007). Retrotransposon silencing by DNA methylation can drive mammalian genomic imprinting. PLoS Genet 3, e55. doi: 10.1371/journal.pgen.0030055

Underhill, D. M., Ozinsky, A., Hajjar, A. M., Stevens, A., Wilson, C. B., Bassetti, M. and Aderem, A (1999). The Toll-like receptor 2 is recruited to macrophage phagosomes and discriminates between pathogens. Nature 401, 811–815. doi: 10.1038/44605.

Underhill, D., M. (2003). Macrophage recognition of zymosan particles. J. Endotoxin Res. 9, 176–180. doi: 10.1179/096805103125001586

Xu, B. and Yang, Z. (2013). PAMLX: a graphical user interface for PAML. Mol. Biol. Evol. 30, 2723–2724. doi:10.1093/molbev/mst179

Xu, H. and Ren, D. (2015). Lysosomal Physiology. Annu. Rev. Physiol. 77, 57–80. doi:10.1146/annurev-physiol-021014-071649

Youngson, N. A., Kocialkowski, S., Peel, N. and Ferguson-Smith, A. C. (2005). A small family of sushi-class retrotransposon-derived genes in mammals and their relation to genomic imprinting. J. Mol. Evol. 61, 481–490. doi:10.1007/s00239-004-0332-0

