## Supplementary material for "Retrovirus-derived *RTL9* plays an important role in innate antifungal immunity in the eutherian brain": figs S1-13

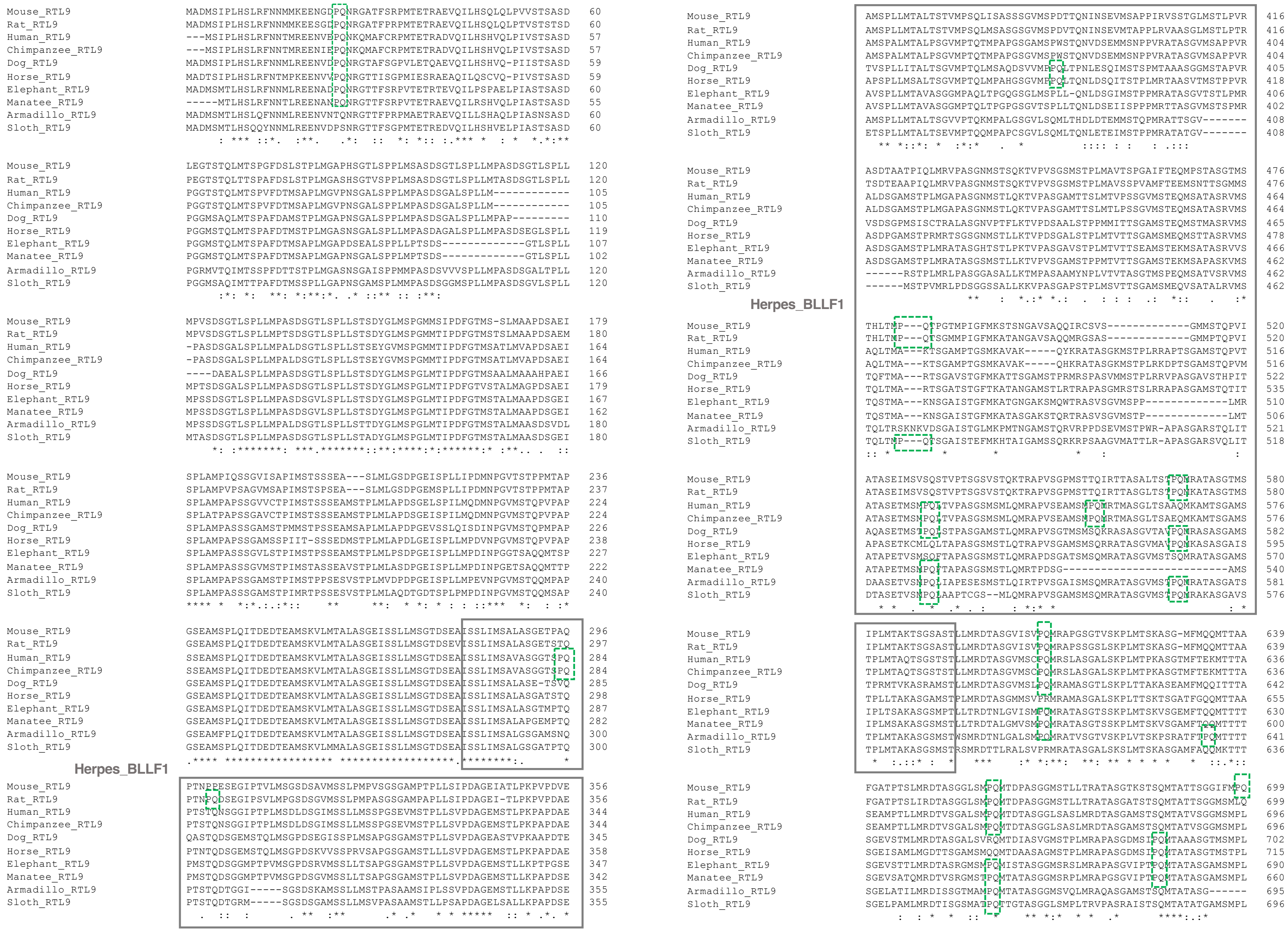

Mouse_RTL9 MADMSIPLHSLRFNNMMKEENGDPQNRGATFSRPMTETRAEVQILHSQLQLPVVSTSASD 60
Rat_RTL9 MADMSIPLHSLRFNNMMKEESGDPQNRGATFPRPMTETRAEVQILHSQLQLPTVSTSTSD 60
Human_RTL9 ---MSIPLHSLRFNNTMREENVEPQNKQMAFCRPMTETRADVQILHSHVQLPIVSTSASD 57
Chimpanzee_RTL9 ---MSIPLHSLRFNNTMREENIEPQNKQMAFCRPMTETRADVQILHSHVQLPIVSTSASD 57
Dog_RTL9 MADMSIPLHSLRFNNMLREENVDPQNRGTAFSGPVLETQAEVQILHSHVQ-PIISTSASD 59
Horse_RTL9 MADTSIPLHSLRFNTMPKEENVVPQNRGTTISGPMIESRAEAQILQSCVQ-PIVSTSASD 59
Elephant_RTL9 MADMSMTLHSLRFNNMLREENADPQNRGTTFSRPVTETRTEVQILPSPAELPIASTSASD 60
Manatee_RTL9 -----MTLHSLRFNNTLREENANPQNRGTTFSRPVTETRAEVQILRSHVQLPIASTSASD 55
Armadillo_RTL9 MADMSMTLHSLQFNNMLREENVNTQNRGTTFPRPMAETRAEVQILLSHAQLPIASNSASD 60
Sloth_RTL9 MADMSMTLHSQQYNNMLREENVDPSNRGTTFSGPMTETREDVQILHSHVELPIASTSASD 60
Mouse_RTL9 LEGTSTQLMTSPGFDSLSTPLMGAPHSGTLSPPLMSASDSGTLSPLLMPASDSGTLSPLL 120
Rat_RTL9 PEGTSTQLTTSPAFDSLSTPLMGASHSGTVSPPLMSASDSGTLSPLLMTASDSGTLSPLL 120
Human_RTL9 PGGTSTQLMTSPVFDTMSAPLMGVPNSGALSPPLMPASDSGALSPLLM------------ 105
Chimpanzee_RTL9 PGGTSTQLMTSPVFDTMSAPLMGVPNSGALSPPLMPASDSGALSPLLM------------ 105
Dog_RTL9 PGGMSAQLMTSPAFDAMSTPLMGAPNSGALSPPLMPASDSGALSPLLMPAP--------- 110
Horse_RTL9 PGGMSTQLMTSPAFDTMSTPLMGASNSGALSPPLLMPASDAGALSPLLMPASDSEGLSPLL 119
Elephant_RTL9 PGGMSTQLMTSPAFDTMSAPLMGAPDSEALSPPLLPTSDS-------------GTLSPLL 107
Manatee_RTL9 PGGMSTQLMTSPAFDTMSAPLMGAPNSGALSPPLMPTSDS-------------GTLSPLL 102
Armadillo_RTL9 PGRMVTQIMTSSPFDTTSTPLMGASNSGAISPPMMPASDSVVVSPLLMPASDSGALTPLL 120
Sloth_RTL9 PGGMSAQIMTTPAFDTMSSPLLGAPNSGAMSPLMMPASDSGGMSPLLMPASDSGVLSPLL 120
Mouse_RTL9 MPVSDSDGTLSPLLMPASDSGTLSPLLSTSDYGLMSPGMMSIPDFGTMS-SLMAAPDSAEI 179
Rat_RTL9 MPVSDSDGTLSPLLMPTSDSGTLSPLLSTSDYGLMSPGMMTIPDFGTMSTSLMAAPDSAEM 180
Human_RTL9 -PASDSGALSPLLMPALDSGTLSPLLSTSEYGVMSPGMMTIPDFGTMSATLMVAPDSAEI 164
Chimpanzee_RTL9 -PASDSGALSPLLMPALDSGTLSPLLSTSEYGVMSPGMMTIPDFGTMSATLMVAPDSAEI 164
Dog_RTL9 ----DAEALSPLLMPASDSGTLSPLLSTSDYGLMSPGLMTIPDFGTMSAALMAAAHPAEI 166
Horse_RTL9 MPTSDSGALSPLLMPASDSGTLSPLLSTSDYGLMSPGLMTIPDFGTVSTALMAGPDSAEI 179
Elephant_RTL9 MPSSDSGTLSPLLMPASDSGVLSPLLSTSDYGLMSPGLMTIPDFGTMSTALMAAPDSGEI 167
Manatee_RTL9 MPSSDSGTLSPLLMPASDSGVLSPLLSTSDYGLMSPGLMTIPDFGTMSTALMAAPDSGEI 162
Armadillo_RTL9 MPSSDSGTLSPLLMPALDSGTLSPLLSTTDYGLMSPGLMTIPDFGTMSTALMAASDSVDL 180
Sloth_RTL9 MTASDSGTLSPLLMPASDSGTLSPLLSTADYGLMSPGLMTIPDFGTMSTALMAASDSGEI 180
Mouse_RTL9 SPLAMPIQSSGVISAPIMSTSSSEA---SLMLGSDPGEISPLLIPDMNPGVTSTPPMTAP 236
Rat_RTL9 SPLAMPVPSAGVMSAPIMSTSPSEA---SLMLGSDPGEMSPLLIPDMNPGVTSTPPMTAP 237
Human_RTL9 SPLAMPAPSSGVVCTPIMSTSSSEAMSTPLMLAPDSGELSPLILMQDMNPGVMSTQPVAP 224
Chimpanzee_RTL9 SPLATPAPSSGAVCTPIMSTSSSEAMSTPLMLAPDSGEISPILMQDMNPGVMSTQPVAP 224
Dog_RTL9 SPLAMPASSSGAMSTPMMSTPSSEAMSAPLMLAPDPGEVSSLQISDINPGVMSTQPMPAP 226
Horse_RTL9 SPLAMPASSSGAMSSPIIT-SSSEDMSTPLMLAPDLGEISPLLMPDVNPGVMSTQPVPAP 238
Elephant_RTL9 SPLAMPASSSGVLSTPIMSTPSSEAMSTPLMLPSDPGEISPLLMPDINPGGTSAQQMTSP 227
Manatee_RTL9 SPLAMPASSSGVMSTPIMSTASSEAVSTPLMLASDPGEISPLLMPDINPGETSAQQMTTP 222
Armadillo_RTL9 SPLAMPASSSGAMSTPIMSTPPSESVSTPLMVDPDPGEISPLLMPEVNPGVMSTQQMPAP 240
Sloth_RTL9 SPLAMPASSSGAMSTPIMRTPSSESVSTPLMLAQDTGDTSPLPMPDINPGVMSTQQMSAP 240
Mouse_RTL9 GSEAMSPLQITDEDTEAMSKVLMTALASGEISSLLMSGTDSEAISSLIMSALASGETPAQ 296
Rat_RTL9 GSEAMSPLQITDEDTEAMSKVLMTALASGEISSLLMSGTDSEVISSLIMSALASGETSTQ 297
Human_RTL9 SSEAMSPLQITDEDTEAMSKVLMTALASGEISSLLMSGTDSEAISSLIMSAVASGGTSPQ 284
Chimpanzee_RTL9 SSEAMSPLQITDEDTEAMSKVLMTALASGEISSLLMSGTDSEAISSLIMSAVASGGTSPQ 284
Dog_RTL9 GSEAMSPLQITDEDTEAMSKVLMTALASGEISSLLMSGTDSEAISSLIMSALASE-TSVQ 285
Horse_RTL9 GSEAMSPLQITDEDTEAMSKVLMTALASGEISSLLMSGTDSEAISSLIMSALASGATSTQ 298
Elephant_RTL9 GSEAMSPLQITDEDTEAMSKVLMTALASGEISSLLMSGTDSEAISSLIMSALASGTMPTQ 287
Manatee_RTL9 GSEAMSPLQITDEDTEAMSKVLMTALASGEISSLLMSGTDSEAISSLIMSALASGEMPTQ 282
Armadillo_RTL9 GSEAMFPLQITDEDTEAMSKVLMTALASGEISSLLMSGTDSEAISSLIMSALGSGAMSNQ 300
Sloth_RTL9 GSEAMSPLQITDEDTEAMSKVLMMALASGEISSLLMSGTDSEAISSLIMSALGSGATPTQ 300
Herpes_BLLF1
Mouse_RTL9 PTNPPESEGIPTVLMSGSDSAVMSSLPMPVSGSGAMPTPLLSIPDAGEIATLPKPVPDVE 356
Rat_RTL9 PTNPQDSEGIPSVLMPGSDSGVMSSLPMPASGSGAMPAPLLSIPDAGEI-TLPKPVPDAE 356
Human_RTL9 PTSTQNSGGIPTPLMSDLDSGIMSSLLMSSPGSEVMSTPLLSVPDAGEMSTLPKPAPDAE 344
Chimpanzee_RTL9 PTSTQNSGGIPTPLMSDLDSGIMSSLLMSSPGSEVMSTPLLSVPDAGEMSTLPKPAPDAE 344
Dog_RTL9 QASTQDSGEMSTQLMSGPDSEGISSPLMSAPGSGAMSTPLLSVPDAGEASTVPKAAPDTE 345
Horse_RTL9 PTNTQDSGEMSTQLMSGPDSKVSSPRVSAPGSGAMSTPLLLSVPDAGEMSTLPKPAPDAE 358
Elephant_RTL9 PMSTQDSGGMPTPVMSGPDSRVMSSLLTSAPGSGAMSTPLLSVPDAGEMSTLLKPTPGSE 347
Manatee_RTL9 PMSTQDSGGMPTPVMSGPDSGVMSSLLTSAPGSGAMSTPLLSVPDAGEMSTLLKPAPDSE 342
Armadillo_RTL9 PTSTQDTGGI-----SGSDSKAMSSLLMSTPASAAMSIPLLSSVPDAGEMSTLLKPAPDSE 355
Sloth_RTL9 PTSTQDTGRM-----SGSDSGAMSSLLMSVPASAAMSTPLLPSAPDAGELSALLKPAPDSE 355
Mouse_RTL9 AMSPLLMTALTSTVMPSQLISASSSGVMSPDTTQNINSEVMSAPPIRVSSTGLMSTLPVR 416
Rat_RTL9 AMSPLLMTALTSTVMPSQLMSASGSGVMSPDVTQNINSEVMTAPPLRVAASGLMSTLPTR 416
Human_RTL9 AMSPALMTALPSGVMPTQTMPAPGSGAMSPWSTQNVDSEMMSNPPVRATASGVMSAPPVR 404
Chimpanzee_RTL9 AMSPALMTALPSGVMPTQTMPAPGSGVMSPWSTQNVDSEMMSNPPVRATASGVMSAPPVR 404
Dog_RTL9 TVSPLLITALTSGVMPTQLMSAQDSVVMPPQLTPNLESQIMSTSPMTAAASSGGMSTAPVR 405
Horse_RTL9 APSPLLMSALTSGVMPTQLMPAHGSGVMPPQLTQNLDSQITSTPLMRTAASVTMSTPPVR 418
Elephant_RTL9 AVSPLLMTAVASGGMPAQLTPGQGSGLMSPLL-QNLDSGIMSTPPMRATASGVTSTLPMR 406
Manatee_RTL9 AVSPLLMTAVASGGMPTQLTPGPGSGVTSPLLTQNLDSEIISPPPMRTTASGVMSTSPMR 402
Armadillo_RTL9 AMSPLLMTALTSGVVPTQKMPALGSGVLSQMLTHDLDTEMMSTQPMRATTSGV------- 408
Sloth_RTL9 ETSPLLMTALTSEVMPTQQMPAPCSGVLSQMLTQNLETEIMSTPPMRATATGV------- 408
Mouse_RTL9 ASDTAATPIQLMRVPASGNMSTSQKTVPVSGSMSTPLMAVTSPGAIFTEQMPSTASGTMS 476
Rat_RTL9 TSDTEAAPIQLMRVPASGNMSTSQKTVPVSGSMSTPLMAVSSPVAMFTEEMSNTTSGMMS 476
Human_RTL9 ALDSGAMSTPLMGAPASGNMSTLQKTVPASGAMTTSLMTVPSSGVMSTEQMSATASRVMS 464
Chimpanzee_RTL9 ALDSGAMSTPLMGAPASGNMSTLQKTVPASGAMTTSLMTLPSSGVMSTEQMSATASRVMS 464
Dog_RTL9 VSDSGPMSISCTRALASGNVPTFLKTVPDSAALSTPPMMITTSGAMSTEQMSTMASRVMS 465
Horse_RTL9 ASDPGAMSTPRMRTSGSGNMSTLLKTVPDSGALSTPLMTVTTSGAMSMEQMSTTASRVMS 478
Elephant_RTL9 ASDSGAMSTPLMRATASGHTSTLPKTVPASGAVSTPLMTVTTSGAMSTEKMSATASRVVS 466
Manatee_RTL9 ASDSGAMSTPLMRATASGSMSTLLKTVPVPSGAMSTPPMTVTTSGAMSTEKMSAPASKVMS 462
Armadillo_RTL9 ------RSTPLMRLPASGGASALLKTMPASAAMYNPLVTVTASGTMSPEQMSATVSRVMS 462
Sloth_RTL9 ------MSTPVMRLPDSGGSSALLKKVPASGAPSTPLMSVTTSGAMSMEQVSATALRVMS 462
Herpes_BLLF1
Mouse_RTL9 THLTMP---QTPGTMPIGFMKSTSNGAVSAQQIRCSVS-------------GMMSTQPVI 520
Rat_RTL9 THLTMP---QTSGMMPIGFMKATANGAVSAQQMRGSAS-------------GMMPTQPVI 520
Human_RTL9 AQLTMA---KTSGAMPTGSMKAVAK-----QYKRATASGKMSTPLRRAPTSGAMSTQPVT 516
Chimpanzee_RTL9 AQLTMA---KTSGAMPTGSMKAVAK-----QHKRATASGKMSTPLRKDPTSGAMSTQPVM 516
Dog_RTL9 TQFTMA---RTSGAVSTGFMKATTSGAMSTPRMRSPASVMMSTPLRRVPASGAVSTHPIT 522
Horse_RTL9 TQLTMA---RTSGAISTGFTKATANGAMSTLRRTAPASGMRSTSLRRAPASGAMSTQTIT 535
Elephant_RTL9 TQSTMA---KNSGAISTGFMKATGNGAKSMQWTRASVSGVMSPP-------------LMR 510
Manatee_RTL9 TQSTMA---KNSGAISTGFMKATASGAKSTQRTRASVSGVMSTP-------------LMT 506
Armadillo_RTL9 TQLTRSKNKVDSGAISTGLMKPMTNGAMSTQRVRPPDSEVMSTPWR-APASGARSTQLIT 521
Sloth_RTL9 TQLTMP---QTSGAISTEFMKHTAIGAMSSQRKRPSAAGVMATTLR-APASGARSVQLIT 518
Mouse_RTL9 ATASEIMSVSQSTVPTSGSVSTQKTRAPVSGPMSTTQIRTTASALTSTPQMRATASGTMS 580
Rat_RTL9 ATASEIMSVSQSTVPTSGSVSTQKTRAPVSGPMSTTQIRTTASGLTSTPQNKATASGTMS 580
Human_RTL9 ATASETMSMPQTVPASGSMSMLQMRAPVSEAMSMPQMRTMASGLTSAAQMKAMTSGAMS 576
Chimpanzee_RTL9 ATASETMSMPQTVPASGSMSMLQMRAPVSEAMSMPQMRTMASGLTSAEQMKAMTSGAMS 576
Dog_RTL9 AQASETMSTPQLSTPASGAMSTLQMRAPVSGTMSMSQKRASASGVTAVPQMRASASGAMS 582
Horse_RTL9 APASETKCMLQLTAPASGSMSTLQTRAPVSGAMSMSQRRATASGVMAVPQMKASASGAIS 595
Elephant_RTL9 ATAPETVSMSQFTAPASGSMSTLQMRAPDSGATSMSQMRATASGVMSTSQMRATASGAMS 570
Manatee_RTL9 ATAPETMSMPQFTAPASGSMSTLQMRTPDSG-------------------------AMS 540
Armadillo_RTL9 DAASETVSMPQLIAPESEMSTLQIRTPVSGAISMSQMRATASGVMSTPQMRATASGATS 581
Sloth_RTL9 DTASETVSMPQLAAPTCGS--MLQMRAPVSGAMSMSQMRATASGVMSTPQMRAKASGAVS 576
Mouse_RTL9 IPLMTAKTSGSASTLLMRDTASGVISVPQMRAPGSGTVSKPLMTSKASG-MFMQQMTTAA 639
Rat_RTL9 IPLMTAKTSGSASTLLMRDTASGVISVPQMRAPSSGSLSKPLMTSKASG-MFMQQMTTAA 639
Human_RTL9 TPLMTAQTSGSTSTLLMRDTASGVMSCPQMRSLASGALSKPLMTPKASGTMFTEKMTTTA 636
Chimpanzee_RTL9 TPLMTAQTSGSTSTLLMRDTASGVMSCPQMRSLASGALSKPLMTPKASGTMFTEKMTTTA 636
Dog_RTL9 TPRMTVKASRAMSTLLMRDTASGVMSLPQMRAMASGTLSKPLTTAKASEAMFMQQITTTA 642
Horse_RTL9 TPLLTAKASGAMSTPLMRDTASGMMSVPRMRAMASGALSKPLTTSKTSGATFGQQMTTAA 655
Elephant_RTL9 IPLTSAKASGSMPTLLTRDTNLGVISMPQMRATASGTSSKPLMTSKVSGEMFTQQMTTTT 630
Manatee_RTL9 IPLMSAKASGSMSTLLTRDTALGMVSMPQMRATASGTSSKPLMTSKVSGAMFTQQMTTTT 600
Armadillo_RTL9 TPLMTAKASGSMSTWSMRDTNLGALSMPQMRATVSGTVSKPLVTSKPSRATFTPQMTTTT 641
Sloth_RTL9 TPLMTAKASGSMSTRSMRDTTLRALSVPRMRATASGALSKSLMTSKASGAMFAQQMKTTT 636
Mouse_RTL9 FGATPTSLMRDTASGGLSMPQMTDPASGGMSTLLTRATASGTKSTSQMTATTSGGIFMPQ 699
Rat_RTL9 FGATPTSLIRDTASGGLSMPQMTDPASGGMSTLLTRATASGATSTSQMTATTSGGMSMLQ 699
Human_RTL9 SEAMPTLLMRDTVSGALSMPQMTDTASGGLSASLMRDTASGAMSTSQMTATVSGGMSMPL 696
Chimpanzee_RTL9 SEAMPTLLMRDTVSGALSMPQMTDTASGGLSASLMRDTASGAMSTSQMTATVSGGMSMPL 696
Dog_RTL9 SGEVSTMLMRDTASGALSVRQMTDIASVGMSTPLMRAPASGDMSIPQMTAAASGTMSMPL 702
Horse_RTL9 SGEISAMLMGDTTSGAMSMQQMTDAASAGMSTPLMRAPASGDMSIPQMTATASGTMSTPL 715
Elephant_RTL9 SGEVSTTLMRDTASRGMSMPQMISTASGGMSRSLMRAPASGVIPTPQMTATASGAMSMPL 690
Manatee_RTL9 SGEVSATQMRDTVSRGMSTPQMTATASGGMSRPLMRAPGSGVIPTPQMTATASGAMSMPL 660
Armadillo_RTL9 SGELATILMRDISSGTMAMPQMTATASGGMSVQLMRAQASGAMSTSQMTATASG------ 695
Sloth_RTL9 SGELPAMLMRDTISGSMATPQTTGTASGGLSMPLTRVPASRAISTSQMTATATGAMSMPL 696


Fig. S1-1

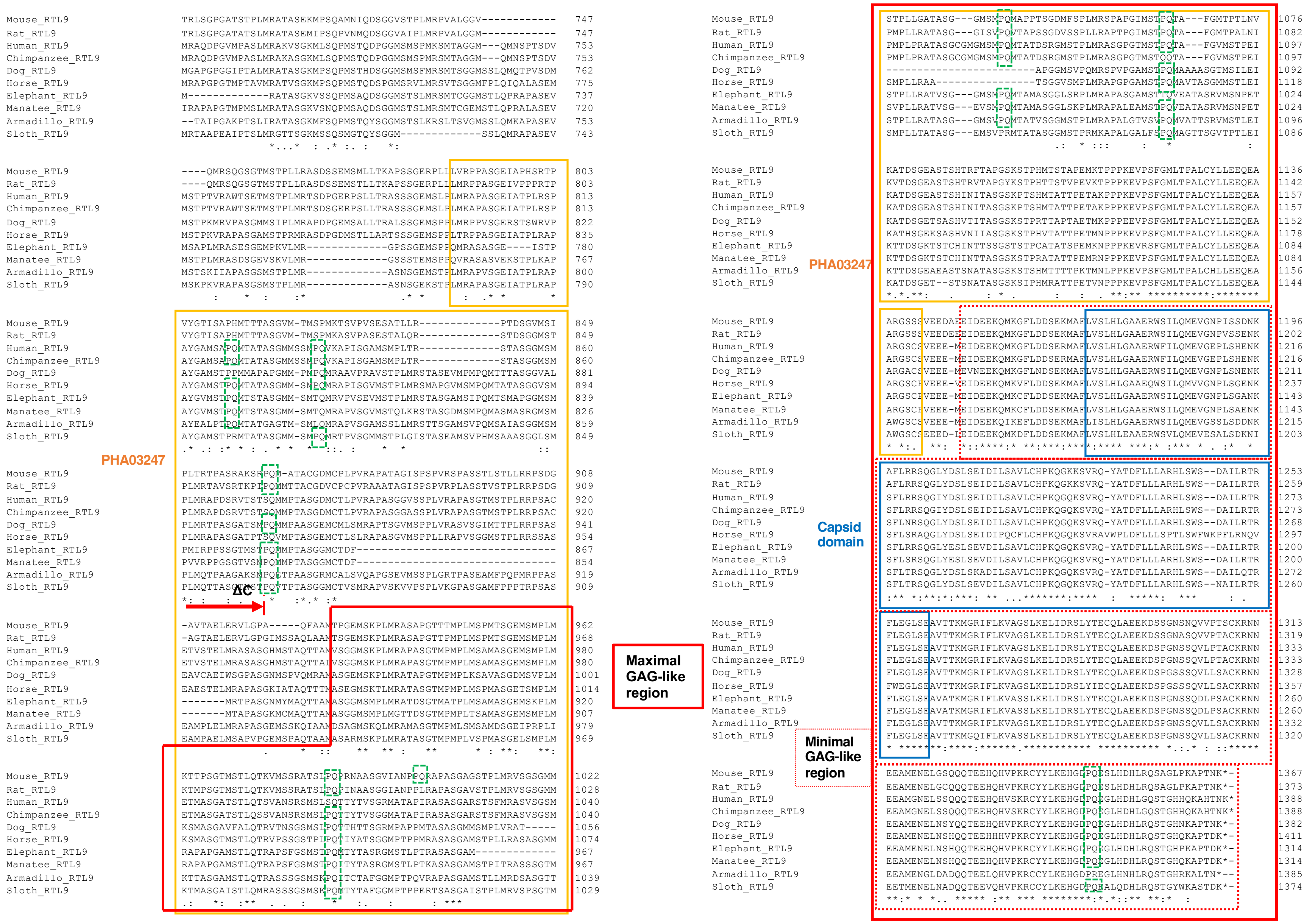
PHA03247
ΔC
Maximal GAG-like region
PHA03247
Capsid domain
Minimal GAG-like region
Mouse_RTL9
Rat_RTL9
Human_RTL9
Chimpanzee_RTL9
Dog_RTL9
Horse_RTL9
Elephant_RTL9
Manatee_RTL9
Armadillo_RTL9
Sloth_RTL9

Fig. S1-2

**Fig. S1 Comparison of the RTL9 protein in the 10 representative eutherian species**

Multiple alignment of the RTL9 protein sequences in 10 eutherian species: the mouse, rat, human, chimpanzee, dog, horse, elephant, manatee, armadillo and sloth. The identical amino acids in the 10 species are depicted with an asterisk (*) below the alignment. Three regions corresponding to the herpes BLLF1 super family, PHA03247 super family and capsid domain of sushi-ichi GAG (Fig. 1A) are shown as gray, orange and blue boxes, respectively. Maximal and minimal GAG-like regions are shown as red boxes and red dashed boxes, respectively. The end of RTL9ΔC, just before the maximal GAG-like region, is indicated by a red arrow with a short vertical line on its head. Details are provided in the Comparative Genome analysis in the Materials and Methods section.

### Conserved domains on [lcl|NM_001040434.2]

View Concise Results

NM_001040434.2 Mus musculus retrotransposon Gag like 9 (Rtl9), mRNA

#### Graphical summary

Zoom to residue level    show extra options »

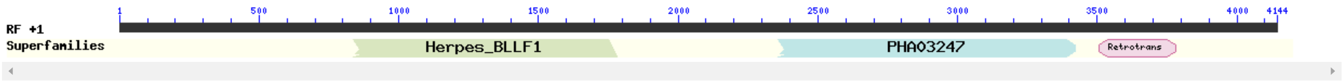


Search for similar domain architectures    Refine search

#### List of domain hits

| Name | Accession | Description | Interval | E-value |
|---|---|---|---|---|
| Retrotrans_gag super family | cl41948 | Retrotransposon gag protein; Gag or Capsid-like proteins from LTR retrotransposons. There is a ... | 3505-3780 | 7.31e-08 |

Retrotransposon gag protein; Gag or Capsid-like proteins from LTR retrotransposons. There is a central motif QGXXEXXXXXFXXLXXH that is common to Retroviridae gag-proteins, but is poorly conserved.

The actual alignment was detected with superfamily member pfam03732:

**Pssm-ID:** 455293 **Cd Length:** 97 **Bit Score:** 51.56 **E-value:** 7.31e-08

```
                              10        20        30        40        50        60        70        80
                      ....*....|....*....|....*....|....*....|....*....|....*....|....*....|....*....|
NM_001040434.2 1169 LVSLHL-GAAERW--SILQMEVGNPISSDN--KAFLRRsqgIYDSLSEIDILSAVLCHPKQGKKSVRQYATDFLLLAR-- 1241
Cdd:pfam03732     2 LAVHSLrGAALTWwkSLVARSIDAFDSWDEIkDAFLKR----FFPSIRKDLLRNELRSLRQGTESVREYVERFKRLARql 77

                              90       100
                      ....*....|....*....|
NM_001040434.2 1242 -HLSWSDAILRTRFLEGLSE 1260
Cdd:pfam03732    78 pHHGRDEEALISAFLRGLRP 97
```

| Name | Accession | Description | Interval | E-value |
|---|---|---|---|---|
| PHA03247 super family | cl33720 | large tegument protein UL36; Provisional | 2359-3423 | 1.61e-06 |

large tegument protein UL36; Provisional

The actual alignment was detected with superfamily member PHA03247:

**Pssm-ID:** 223021 [Multi-domain] **Cd Length:** 3151 **Bit Score:** 53.02 **E-value:** 1.61e-06

```
                              10        20        30        40        50        60        70        80
                      ....*....|....*....|....*....|....*....|....*....|....*....|....*....|....*....|
NM_001040434.2  787 LVRPPASGEIAPHSRTPVYGTISAPHMTTTASGVMTMSPMKTSVPVSESATLLRPTDSGVMSIPLTRTPASRAKSRPQMA 866
Cdd:PHA03247   2698 LADPPPPPPTPEPAPHALVSATPLPPGPAAARQASPALPAAPAPPAVPAGPATPGGPARPARPPTTAGPPAPAPPAAPAA 2777

                              90       100       110       120       130       140       150       160
                      ....*....|....*....|....*....|....*....|....*....|....*....|....*....|....*....|
NM_001040434.2  867 TAcgdmcpIPVRAPATAGISPSPVRSPASStlstllrrPSDGAVTAELERVLGPAQFAAMTPGEMSKPLMRA--SAPGTT 944
Cdd:PHA03247   2778 GP-------PRRLTRPAVASLSESRESLPS--------PWDPADPPAAVLAPAAALPPAASPAGPLPPPTSAqpTAPPPP 2842

                             170       180       190       200       210       220       230       240
                      ....*....|....*....|....*....|....*....|....*....|....*....|....*....|....*....|
NM_001040434.2  945 TMPLMSPMTSGEMSMP---LMKTTPSGTMSTLQTKVMSSRATSLPQPrnaasgvianPPQRAPASGAGSTPLMRVSGSGM 1021
Cdd:PHA03247   2843 PGPPPPSLPLGGSVAPggdVRRRPPSRSPAAKPAAPARPPVRRLARP----------AVSRSTESFALPPDQPERPPQPQ 2912

                             250       260       270       280       290       300       310       320
                      ....*....|....*....|....*....|....*....|....*....|....*....|....*....|....*....|
NM_001040434.2 1022 MSTPLLGATASGGMSMPQMAPPTSGDMFSPLMRSPAPGIMSTPQtafGMTPTLNVKATDSGEASTSHTRFtapgskSTPH 1101
Cdd:PHA03247   2913 APPPPQPQPQPPPPPQPQPPPPPPPRPQPPLAPTTDPAGAGEPS---GAVPQPWLGALVPGRVAVPRFRV------PQPA 2983

                             330       340       350       360
                      ....*....|....*....|....*....|....*....|
NM_001040434.2 1102 MTSTAPEMKTPPPKEVPSFGMLTPALCYLLEEQEAARGSS 1141
Cdd:PHA03247   2984 PSREAPASSTPPLTGHSLSRVSSWASSLALHEETDPPPVS 3023
```

**Fig. S2-1**

Herpes_BLLF1 super family cl37540 Herpes virus major outer envelope glycoprotein (BLLF1); This family consists of the BLLF1 ... 838-1782 5.67e-03

Herpes virus major outer envelope glycoprotein (BLLF1); This family consists of the BLLF1 viral late glycoprotein, also termed gp350/220. It is the most abundantly expressed glycoprotein in the viral envelope of the Herpesviruses and is the major antigen responsible for stimulating the production of neutralising antibodies in vivo.

The actual alignment was detected with superfamily member pfam05109:

**Pssm-ID:** 282904 [Multi-domain] **Cd Length:** 886 **Bit Score:** 41.06 **E-value:** 5.67e-03

```
                           10        20        30        40        50        60        70        80
NM_001040434.2  280 ISSLIMSALASGETPAQPTNPPESEGIPTVL--MSGSDSAVMSSLPMPVSGSGAM--PTPLLSIPDAGEIA---TLPKPV 352
Cdd:pfam05109   473 VTSPTPAGTTSGASPVTPSPSPRDNGTESKApdMTSPTSAVTTPTPNATSPTPAVttPTPNATSPTLGKTSptsAVTTPT 552

                           90       100       110       120       130       140       150       160
NM_001040434.2  353 PDVEAMSPLLMTALTSTVMPSQLISASSSGVMSPdtTQNINSEVMSAPPIRVSST----GLMSTLPVRASDTAAtpiqlm 428
Cdd:pfam05109   553 PNATSPTPAVTTPTPNATIPTLGKTSPTSAVTTP--TPNATSPTVGETSPQANTTnhtlGGTSSTPVVTSPPKN------ 624

                          170       180       190       200       210       220       230       240
NM_001040434.2  429 rvpASGNMSTSQKTVPVSGSMSTPLmavtSPGAIFTEQMPSTASGTMSThltMPQTPGTMPIGFMKSTSNGAVSAQQIRC 508
Cdd:pfam05109   625 ---ATSAVTTGQHNITSSSTSSMSL----RPSSISETLSPSTSDNSTSH---MPLLTSAHPTGGENITQVTPASTSTHHV 694

                          250       260       270       280       290       300       310       320
NM_001040434.2  509 SVSGMMSTQPVIATASEIMSVSQSTVPTSGSVSTQKTRAPVSGPMSTTQIRTTASALTSTPQMRATASGTMSIPLMTAKT 588
Cdd:pfam05109   695 STSSPAPRPGTTSQASGPGNSSTSTKPGEVNVTKGTPPKNATSPQAPSGQKTAVPTVTSTGGKANSTTGGKHTTGHGART 774

NM_001040434.2  589 SGSAST 594
Cdd:pfam05109   775 STEPTT 780
```

### Fig. S2 The result of a conserved domain search using NCBI CDD

Three domains, the retrotransposon gag protein, large tegument protein UL36 and herpes virus major outer envelope glycoprotein (BLLF1), are listed in this order. The identical amino acids in the superfamily motifs and RTL9 are depicted in red.

Fig. S2-2

```
Mouse RTL9      1 ISSLIMSALASGETPAQPTNPPESEGIPTVL--MSGSDSAVMSSLPMPVSGSGAM--PTPLLSIPDAGEIA---TLPKPV     80
                  ::|...:...||.:|..|:..|...|..:..  |:...|||.:..|...|.:.|:  |||..:.|..|:.:   .:..|.
pfam05109       1 VTSPTPAGTTSGASPVTPSPSPRDNGTESKApdMTSPTSAVTTPTPNATSPTPAVttPTPNATSPTLGKTSptsAVTTPT     80

Mouse RTL9     81 PDVEAMSPLLMTALTSTVMPSQLISASSSGVMSPdtTQNINSEVMSAPPIRVSST----GLMSTLPVRASDTAAtpiqlm    160
                  |:..:.:|.:.|...:..:|:...::.:|.|.:|..|.|..|..:.......::|    |..|:.||..|....
pfam05109      81 PNATSPTPAVTTPTPNATIPTLGKTSPTSAVTTP--TPNATSPTVGETSPQANTTnhtlGGTSSTPVVTSPPKN------    160

Mouse RTL9    161 rvpASGNMSTSQKTVPVSGSMSTPLmavtSPGAIFTEQMPSTASGTMSThltMPQTPGTMPIGFMKSTSNGAVSAQQIRC    240
                  ...|:..::|.|..:..|.:.|..|.....|.:|.....|||:..:.|..  ||......|.|....|.....|......
pfam05109     161 ---ATSAVTTGQHNITSSSTSSMSL----RPSSISETLSPSTSDNSTSH---MPLLTSAHPTGGENITQVTPASTSTHHV    240

Mouse RTL9    241 SVSGMMSTQPVIATASEIMSVSQSTVPTSGSVSTQKTRAPVSGPMSTTQIRTTASALTSTPQMRATASGTMSIPLMTAKT    320
                  |.|.........:.||...:.|.||.|...:|:........:.|.:.:..:|....:|||.....:.:|........|:|
pfam05109     241 STSSPAPRPGTTSQASGPGNSSTSTKPGEVNVTKGTPPKNATSPQAPSGQKTAVPTVTSTGGKANSTTGGKHTTGHGART    320

Mouse RTL9    321 SGSAST    326
                  |...:|
pfam05109     321 STEPTT    326
```

Herpes_BLLF1 super family
identity 73/326 = 22.4%
Similarity 125/326 = 38.3%

**Fig. S3 Pairwise alignment and identity between pfam05109 (BLLF1) and mouse RTL9**

The N-terminal part of mouse RTL9 (280 aa – 594 aa) possessing homology to herpes BLLF1 super family motif was re-analyzed using EMBOS Water program and EMBOSS needle program as described in Materials and Methods.

```
Mouse RTL9    1 LVRPPASGEIAPHSRTPVYGTISAPHMTTTASGVMTMSPMKTSVPVSESATLLRPTDSGVMSIPLTRTPASRAKSRPQMA     80
                |..||........:...:......|.....|.......|...:.|...:........:.....|.|..|.:.|......|
PHA03247      1 LADPPPPPPTPEPAPHALVSATPLPPGPAAARQASPALPAAPAPPAVPAGPATPGGPARPARPPTTAGPPAPAPPAAPAA     80

Mouse RTL9   81 TAcgdmcplPVRAPATAGISPSPVRSPASStlstllrrPSDGAVTAELERVLGPAQFAAMTPGEMSKPLMRA--SAPGTT    160
                ..       |.|....|..|.|..|....|        |.|.|...........|...|.:|.....|...|  :||...
PHA03247     81 GP-------PRRLTRPAVASLSESRESLPS--------PWDPADPPAAVLAPAAALPPAASPAGPLPPPTSAqpTAPPPP    160

Mouse RTL9  161 TMPLMSPMTSGEMSMP---LMKTTPSGTMSTLQTKVMSSRATSLPQPrnaasgvianPPQRAPASGAGSTPLMRVSGSGM    240
                ..|....:..|....|   :.:..||.:.:.............|.:|          ...|:..|.|.............
PHA03247    161 PGPPPPSLPLGGSVAPggdVRRRPPSRSPAAKPAAPARPPVRRLARP----------AVSRSTESFALPPDQPERPPQPQ    240

Mouse RTL9  241 MSTPLLGATASGGMSMPQMAPPTSGDMFSPLMRSPAPGIMSTPQtafGMTPTLNVKATDSGEASTSHTRFtapgskSTPH    320
                ...|............||..||.......||..:..|.....|.   |..|...:.|...|..:....|.      ..|.
PHA03247    241 APPPPQPQPQPPPPPQPQPPPPPPPRPQPPLAPTTDPAGAGEPS---GAVPQPWLGALVPGRVAVPRFRV------PQPA    320

Mouse RTL9  321 MTSTAPEMKTPPPKEVPSFGMLTPALCYLLEEQEAARGSS    360
                .:..||...|||........:.:.|....|.|:......|
PHA03247    321 PSREAPASSTPPLTGHSLSRVSSWASSLALHEETDPPPVS    360
```

PHA03247 super family
identity 63/360 = 17.5%
similarity 85/360 = 23.6%

**Fig. S4 Pairwise alignment and identity between PHA03247 family and mouse RTL9 protein**

The middle of mouse RTL9 (787 aa –1141 aa) possessing homology to the PHA03247 super family motif was re-analyzed using the EMBOSS Water program and EMBOSS needle program.

```
GAG            131  ELLFRHQPSRFVSDEAKVGFITSLLADKALSWAIAAVDLDPRLSSDYSAF   180
                    |:....|...|:.|..|:.|:.||....|..|:|..:::...:|||..||
Mouse RTL9    1148  EIDEEKQMKGFLDDSEKMAFLVSLHLGAAERWSILQMEVGNPISSDNKAF  1197

GAG            181  RREFKAVFEHPTYGEDAASRLLALQQGSRSVAEYTLEFRILAAESRWGET   230
                    .|..:.:::..:..:..::.|...:||.:||.:|..:|.:||....|.:.
Mouse RTL9    1198  LRRSQGLYDSLSEIDILSAVLCHPKQGKKSVRQYATDFLLLARHLSWSDA  1247

GAG            231  ALRSAYRRGLSEAIKDLIVR---DRPSSLNELITLSLQMDERLRERRQER   277
                    .||:.:..|||||:...:.|   ....||.|||..||..:.:|.|.:.
Mouse RTL9    1248  ILRTRFLEGLSEAVTTKMGRIFLKVAGSLKELIDRSLYTECQLAEEKD--  1295

GAG            278  AQRAGGSTRQLSHRTSSAPDFSLTSTAAPPPHILLQSPAHPSPRVGEEPM   327
                       :.|::.|:      .|    ||                ..|..||.|
Mouse RTL9    1296  ---SSGNSNQV------VP----TS----------------CKRNNEEAM  1316

GAG            328  --QIGRSRLSRQEREQRLR-DQLCLYCGNNG--------HFIQACPVRPK   366
                      ::|    |:|:.|:... .:.|.|...:|        |..|:..: ||
Mouse RTL9    1317  ENELG----SQQQTEEHQHVPKRCYYLKEHGDPQESLHDHLRQSAGL-PK  1362

GAG            367  GPAHQ    371
                    .|.::
Mouse RTL9    1363  APTNK   1367
```

Red: Amino Acids encoded in the exon 2

Light blue: CCHC RNA-binding motif

Sushi-ichi GAG
identity 65/255 = 25.5%
similarity 110/255 = 43.1%

**Fig. S5 Pairwise alignment and identity between sushi-ichi GAG and mouse RTL9 (1)**

A conserved domain search using NCBI CDD indicated that the C-terminal portion of mouse RTL9 (1169 aa –1260 aa) possesses homology to the Capsid motif of sushi-ichi retrotransposon GAG. However, the regions before and after the Capsid motif (1148 aa-1168 aa and 1261 aa-1367 aa) also exhibit homology to GAG (**Minimal GAG-like region**). The AAs in *Rtl9* exon 2 are shown in red. The AAs of the CCHC motif of GAG are shown in light blue.

```
GAG            6 TPGQ-------PGGPRTPTLP--SPLERRVEAHSAQLSSLQSELTKAFPT     46
                 |||:       ...|.|.|:|  ||:.      |.::|   ..|.|..|:
Mouse RTL9   927 TPGEMSKPLMRASAPGTTTMPLMSPMT------SGEMS---MPLMKTTPS    967

GAG           47 IQGEISELQSSSQTTSSTLSALSNQMSAMATVLASIIQK-----LGSDP-     90
                   |.:|.||  ::..||..::|....:|.:.|:|:..|:     .||.|
Mouse RTL9   968 --GTMSTLQ--TKVMSSRATSLPQPRNAASGVIANPPQRAPASGAGSTPL   1013

GAG           91 --------------GGAAPSEPSLP-LSPRAEPNLASP------------    113
                               |..|....|:| ::|....::.||
Mouse RTL9  1014 MRVSGSGMMSTPLLGATASGGMSMPQMAPPTSGDMFSPLMRSPAPGIMST   1063

GAG          114 --RVFG------------GDFDL---------GKGFLHQ-----------    129
                   ..||            |:...         .|...|.
Mouse RTL9  1064 PQTAFGMTPTLNVKATDSGEASTSHTRFTAPGSKSTPHMTSTAPEMKTPP   1113

GAG          130 --------------CELLFRHQPSR--------------------FVSDE    145
                               |.||...:.:|                    |:.|.
Mouse RTL9  1114 PKEVPSFGMLTPALCYLLEEQEAARGSSSVEEDAEEIDEEKQMKGFLDDS   1163

GAG          146 AKVGFITSLLADKALSWAIAAVDLDPRLSSDYSAFRREFKAVFEHPTYGE    195
                 .|:.|:.||....|..|:|..:::...:|||..||.|..:.:::..:..:
Mouse RTL9  1164 EKMAFLVSLHLGAAERWSILQMEVGNPISSDNKAFLRRSQGLYDSLSEID   1213

GAG          196 DAASRLLALQQGSRSVAEYTLEFRILAAESRWGETALRSAYRRGLSEAIK    245
                 ..::.|...:||.:||.:|..:|.:||....|.:..||:.:..|||||:.
Mouse RTL9  1214 ILSAVLCHPKQGKKSVRQYATDFLLLARHLSWSDAILRTRFLEGLSEAVT   1263

GAG          246 DLIVR---DRPSSLNELITLSLQMDERLRERRQERAQRAGGSTRQLSHRT    292
                 ..:.|   ....||.|||..||..:.:|.|.:.     :.|::.|:
Mouse RTL9  1264 TKMGRIFLKVAGSLKELIDRSLYTECQLAEEKD-----SSGNSNQV----   1304

GAG          293 SSAPDFSLTSTAAPPPHILLQSPAHPSPRVGEEPM--QIGRSRLSRQERE    340
                   .|    ||                ..|..||.|  ::|    |:|:.|
Mouse RTL9  1305 --VP----TS----------------CKRNNEEAMENELG----SQQQTE   1328

GAG          341 QRLR-DQLCLYCGNNG--------HFIQACPVRPKGPAHQ    371
                 :... .:.|.|...:|        |..|:..: ||.|.::
Mouse RTL9  1329 EHQHVPKRCYYLKEHGDPQESLHDHLRQSAGL-PKAPTNK   1367
```

Sushi-ichi GAG
identity 108/490 = 21.8%
similarity 175/490 = 35.7%

**Fig. S6 Pairwise alignment and identity between sushi-ichi GAG and mouse RTL9 (2)**

Entire sushi-ichi GAG exhibits homology to mouse RTL9 (927 aa-1357 aa, **Maximal GAG-like region**).

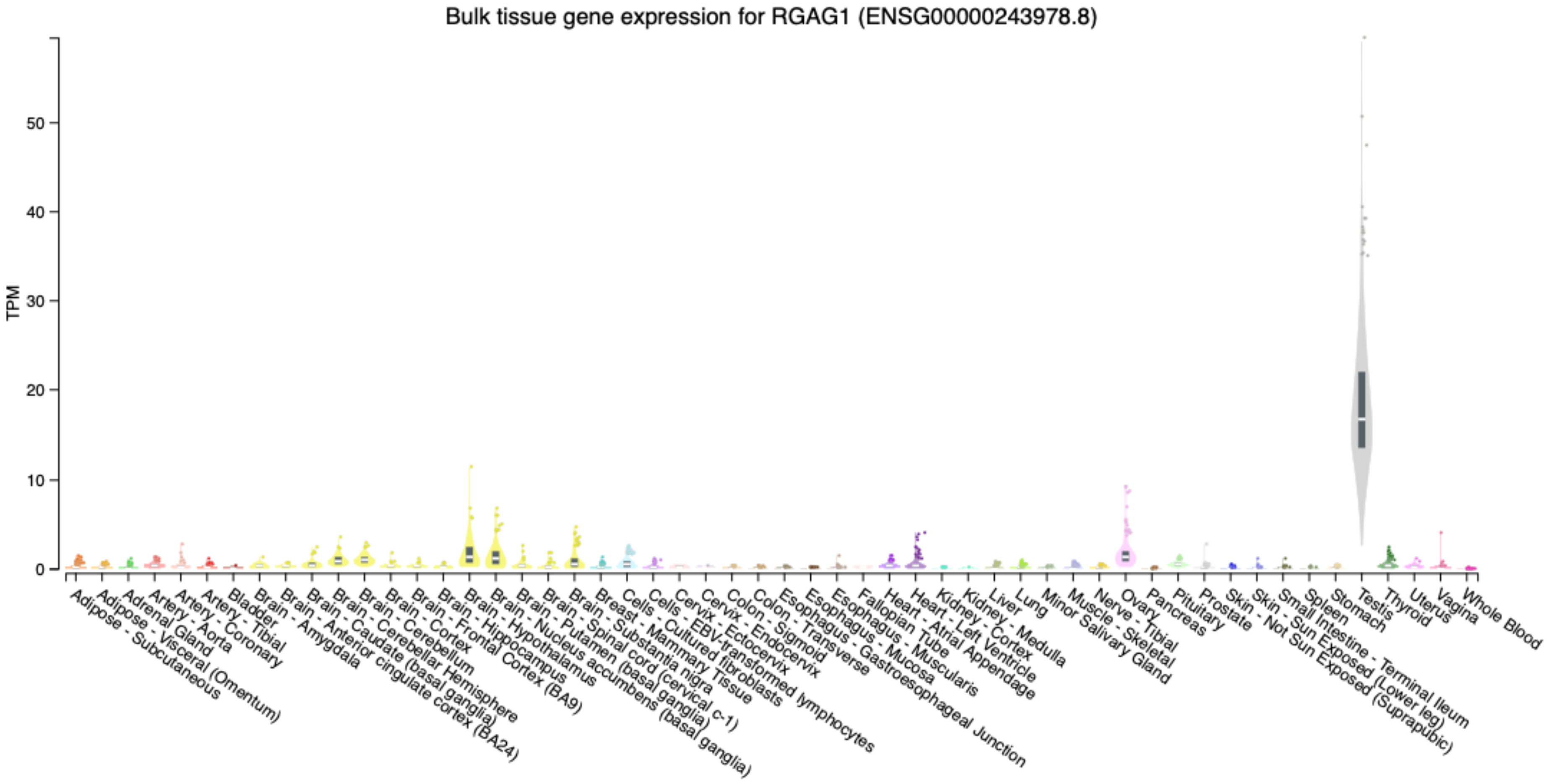


**Fig. S7** ***RTL9*** **(*RGAG1*) expression in human tissues and organs**

See the original data at https://gtexportal.org/home/gene/RGAG1

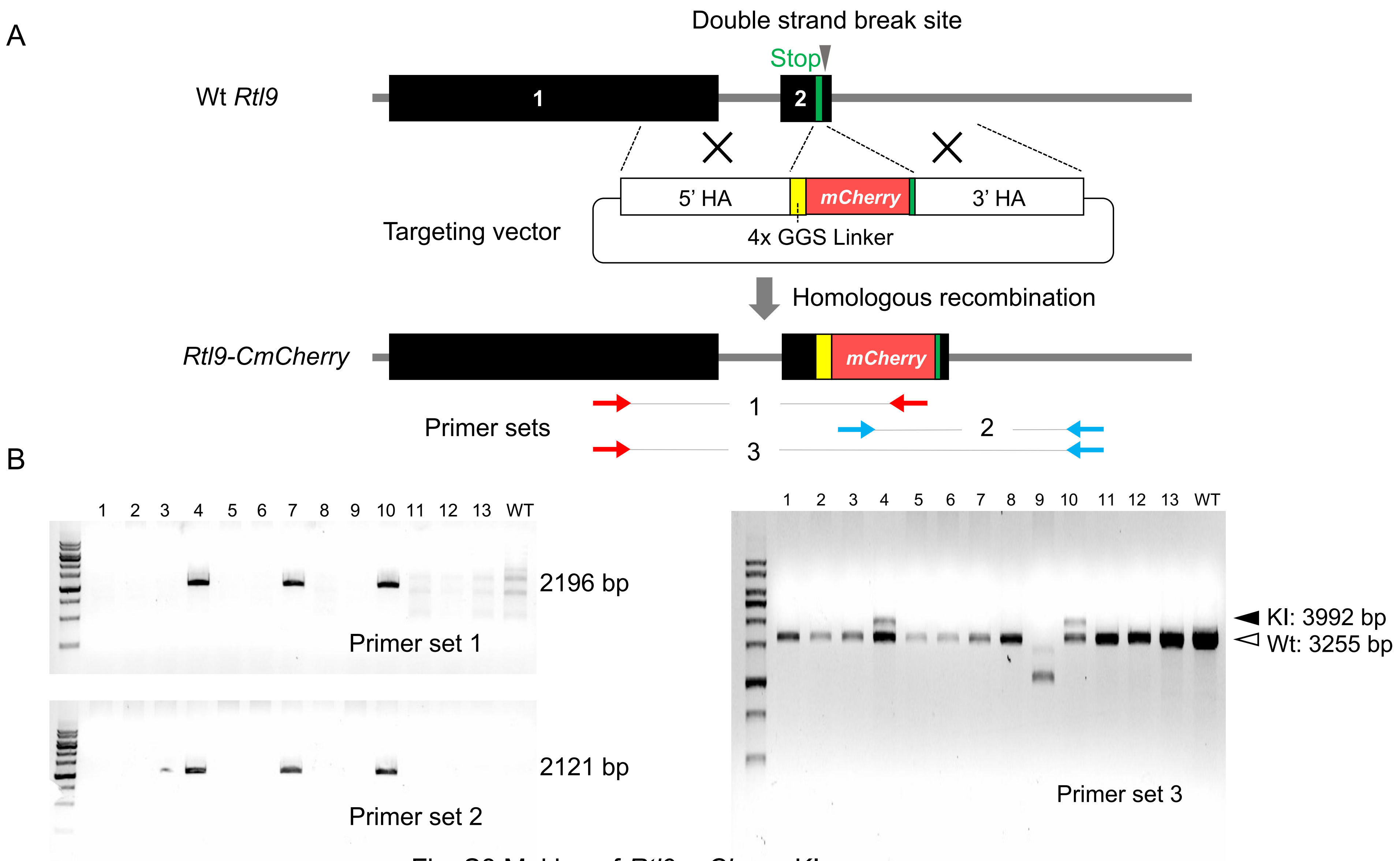


Fig. S8 Making of *Rtl9-mCherry* KI mouse

### Fig. S8 Generation of the *Rtl9CmC* KI mouse

**A)** Schematic representation of the generation of the *Rtl9CmC* KI mouse. *Rtl9* exon 1 and 2 are numbered and shown in black. The stop codon is shown in green. The genomic locus near the stop codon in exon2 of *Rtl9* was cut out using the CRISPR/Cas system and homologous recombination was induced between the Wt *Rtl9* locus and a targeting vector containing 5' and 3' homology arms (5' HA and 3' HA), a 4x GGS linker and the mCherry coding sequence. The reconstructed *Rtl9CmC* has mCherry at the C-terminus. The position of the PCR primer sets for genotyping is indicated. **B)** The results of genotyping 13 founder (F0) mice. Genomic PCR was performed using the primer sets 1 to 3 shown in A. The number (#) of F0 mice is indicated above the gel images. Wild type (WT) mouse genome was used as a control. Based on the genotyping PCR data, the offspring of F0 mouse #4 and #10 were used for further experiments.

PCR primer sets for genotyping of KI and KO mice

| Mutant | Primer set name | Direction | Sequence |
|---|---|---|---|
| *Rtl9-CmCherry* | Primer set 1 | F | GGAATGATGTCCACGCCACTA |
| | | R | CTTCAGCTTCAGCCTCTGCT |
| | primer set 2 | F | CCTGTCCCCTCAGTTCATGT |
| | | R | CCTAGACTATTGGACCAGAGG |
| | Primer set 3 | F | GGAATGATGTCCACGCCACTA |
| | | R | CCTAGACTATTGGACCAGAGG |
| *Rtl9*-KO | Primer set 4 | F | GAGAACACCAGCTTCTAGAGC |
| | | R | GGGAGTTCAGAACCTCATACAC |

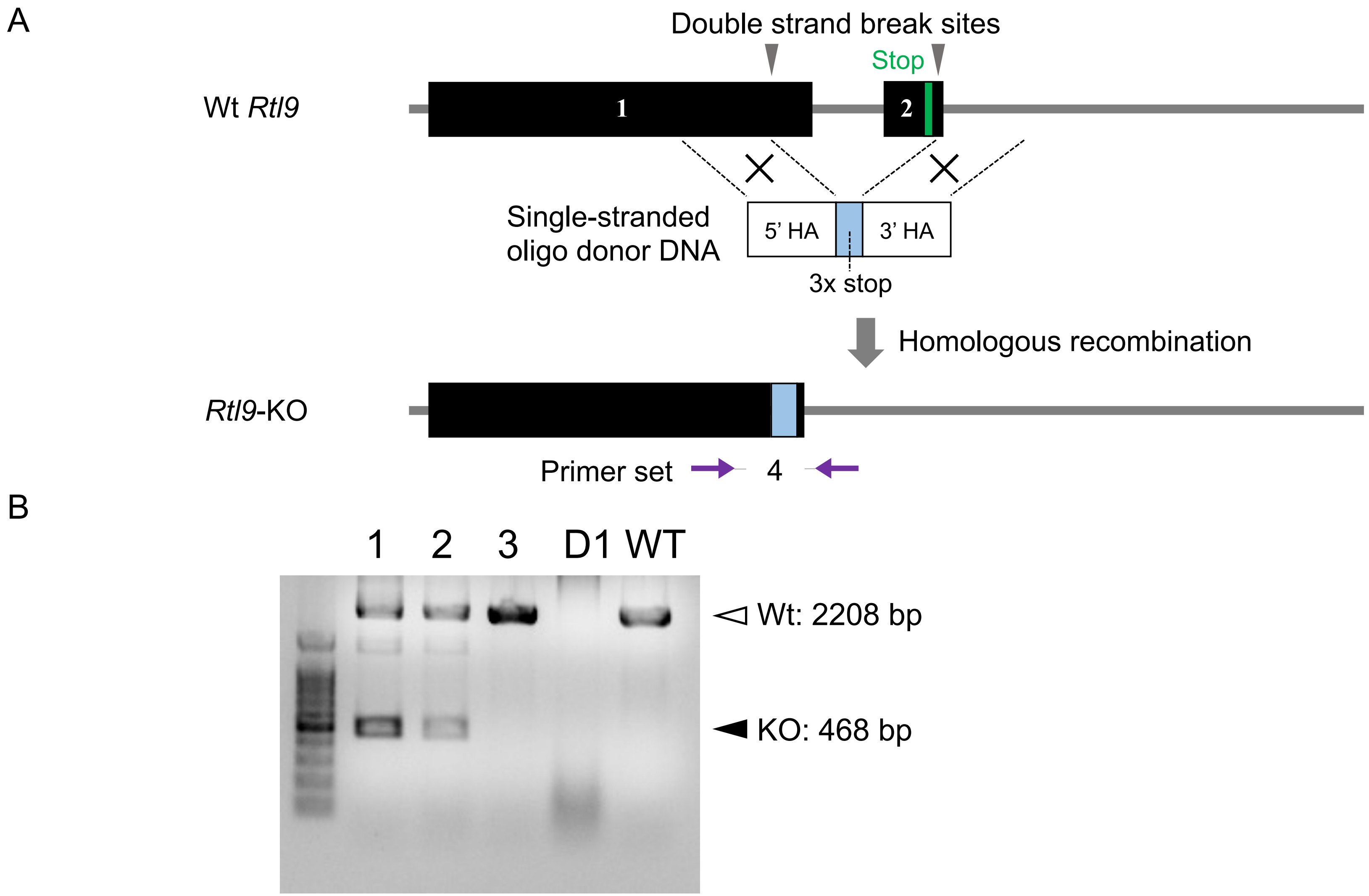


Fig. S9-1 Making of *Rtl9 KO* mouse

C

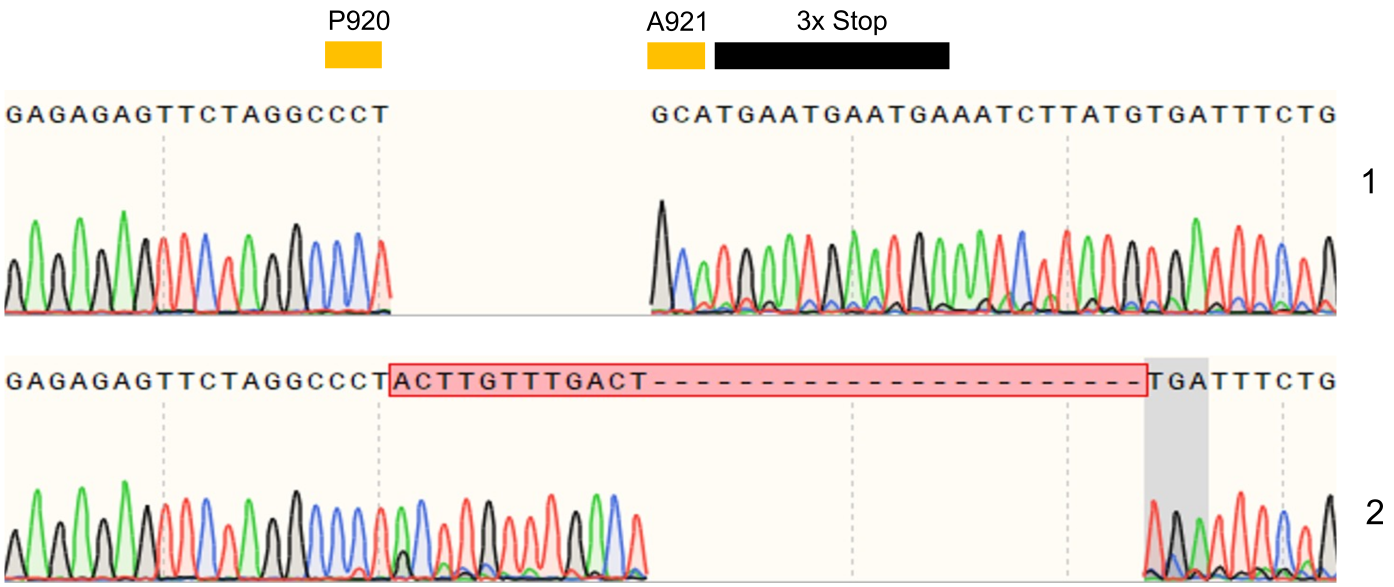


Fig. S9-2 Making of *Rtl9 KO* mouse

### Fig. S9 Making of *Rtl9* KO mouse

**A)** Schematic representation of the generation of the *Rtl9* KO mouse. *Rtl9* exons 1 and 2 are numbered and shown in black. The stop codon is shown in green. The genomic locus encoding alanine 921 (A921) and downstream of the *Rtl9* stop codon was cut using the CRISPR/Cas system, and homologous recombination was induced between the Wt *Rtl9* locus and single-stranded oligo donor DNA containing a sequence of 3 stop codons (3x stop) and 5' and 3' homology arms (5' HA and 3' HA). In the reconstructed *Rtl9*-KO allele, the genomic region from glutamine 922 to the stop codon of *Rtl9* was deleted and replaced by a 3x stop. The position of the PCR primer set used for genotyping is indicated. **B)** Results of genotyping of 3 surviving and 1 dead (D1) F0 mice. Genomic PCR was performed with the primer set 4 shown in A. The number (#) of F0 mice, including D1, is indicated above the gel image. Wild type (WT) mouse genome was used as a control. **C)** Illustrated results of Sanger sequencing using the founder (F0) mice #1 and 2 shown in B. In F0 mouse #1 (top), a 3x stop was inserted at the expected genomic position just after A921 of *Rtl9*. In #2 (bottom), 12 extra bases (highlighted in red) were unexpectedly inserted just before a stop codon (highlighted in grey). Based on the Sanger sequencing data, the offspring of F0 mouse #1 were used for further experiments. The PCR primer sets for genotyping the KO mice are presented in the Fig. S6 legend.

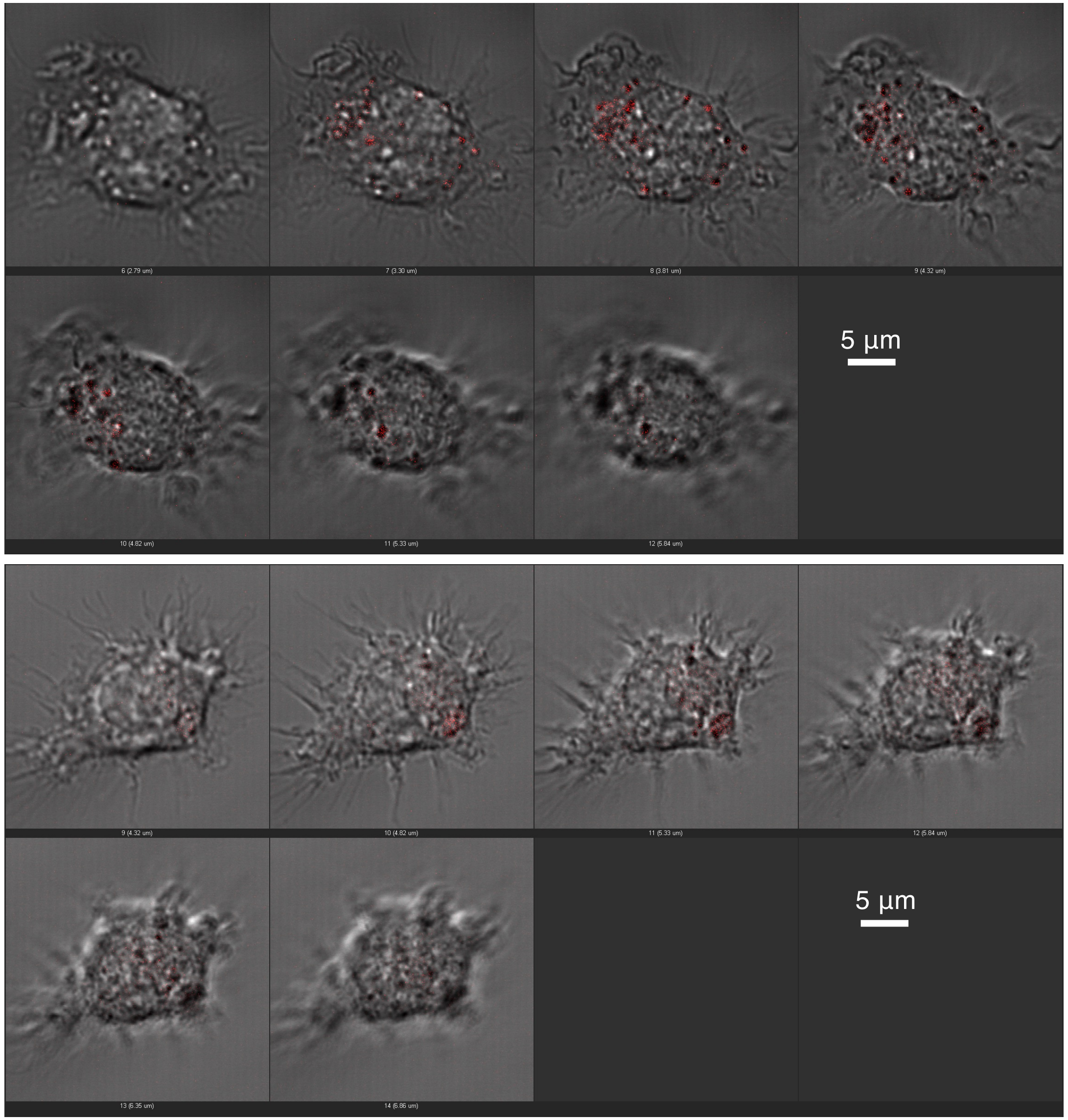


**Fig. S10 mCherry signals in isolated microglia isolated from *Rtl9CmC* KI brain**

Microglia were isolated from P0 neonatal brain and cultured for 10 days. The microglial cells were collected from the cultured Petri dishes (mixed glia cultures) by tapping (Lian et al., 2016; Irie et al., 2022).

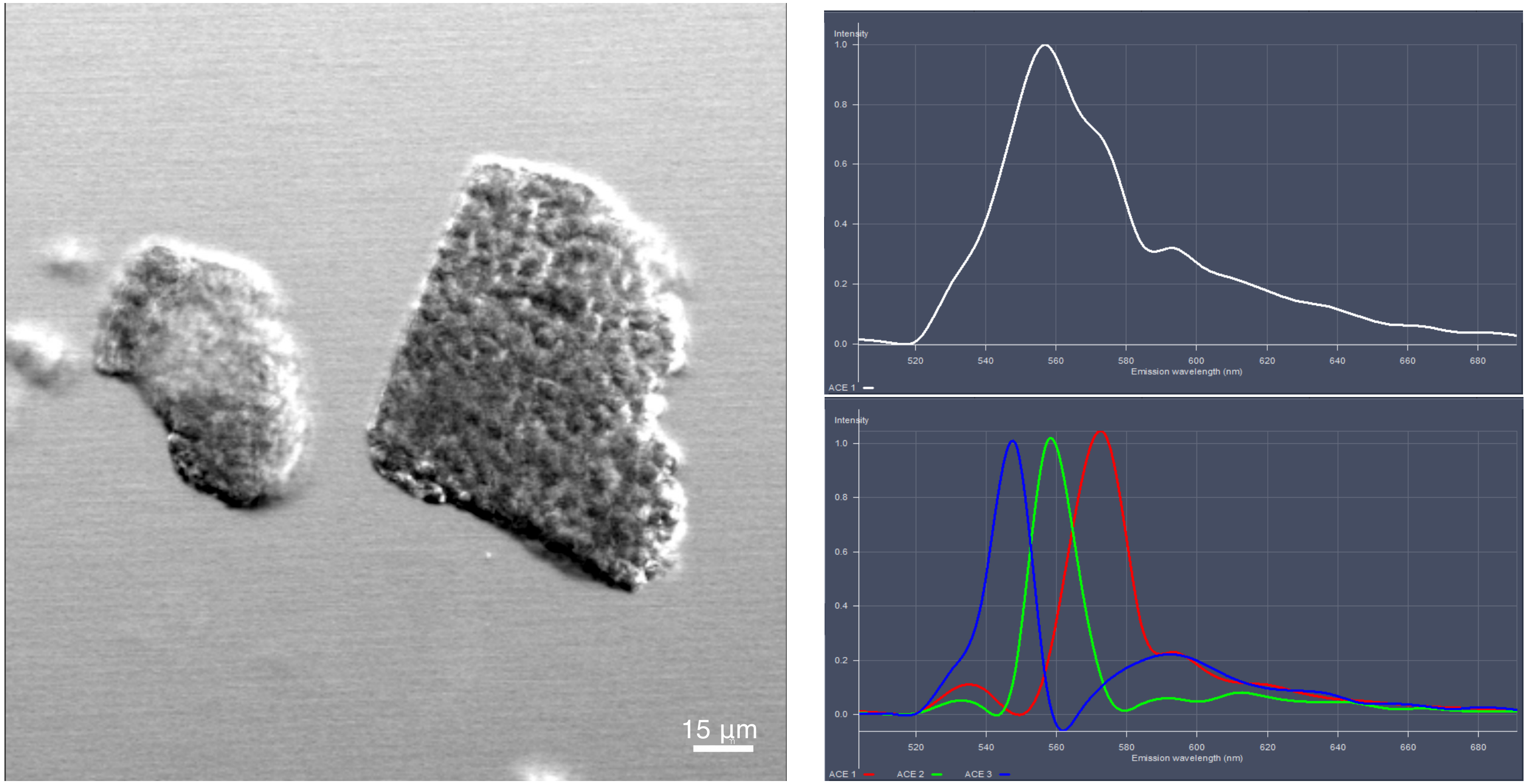


**Fig. S11 The autofluorescence spectrum of Zymosan**

Left: Transmission image. Right: Autofluorescence spectrum. Single (top) and/or the strongest signal (Maximum peak emission fluorescence wavelength: 570 nm) among 3 representative signals were used to detect zymosan.

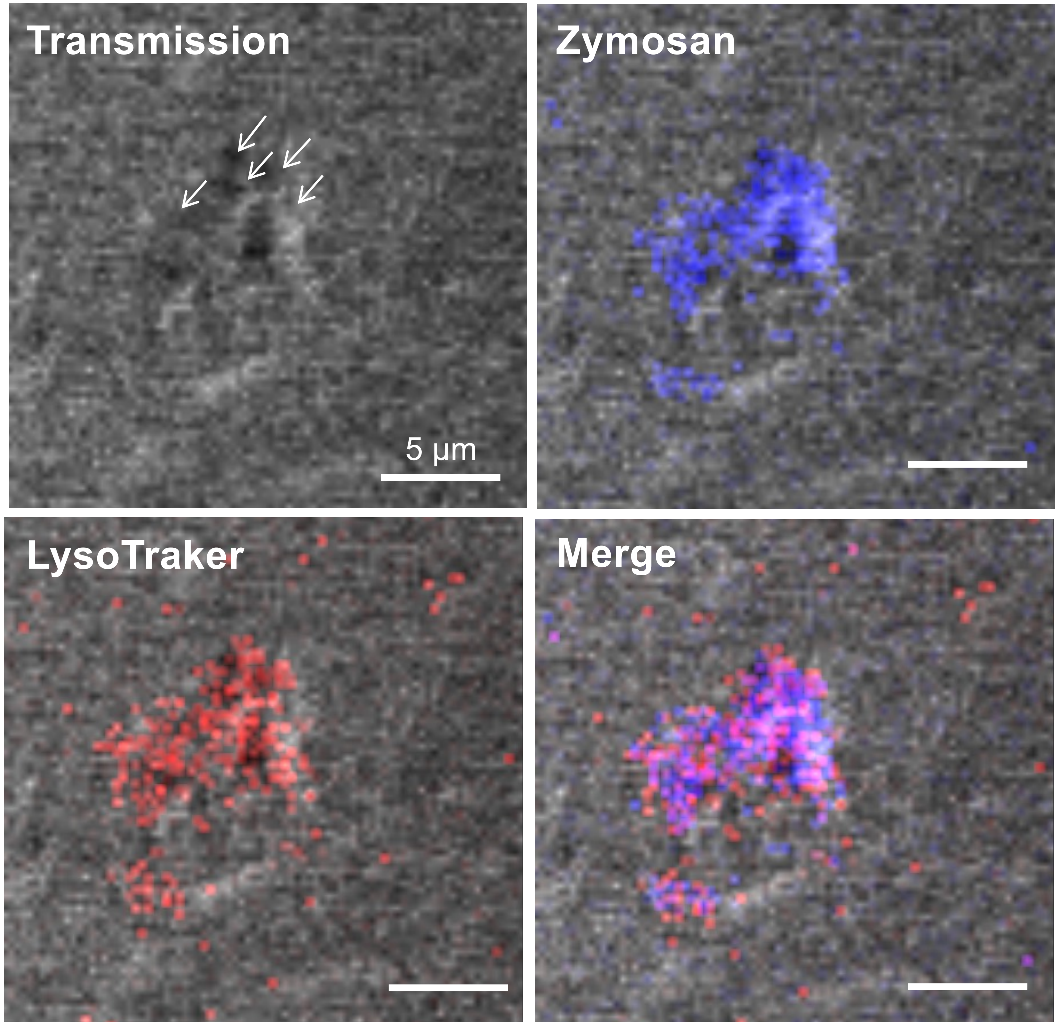


**Fig. S12 Zymosan is visible in the transmission image**

Zymosan incorporated in lysosomes in the *Rtl9* KO brain. Top, left: A transmission image. Arrows indicate zymosan aggregates. Top, right: zymosan autofluorescence (light blue). Bottom, left: The LysoTracker signal (red). Bottom, right: A merged image. Zymosan is distinguishable by its shape in the transmission image(s) in addition to its specific fluorescence signal. The same samples as in Fig. 3E were analyzed.

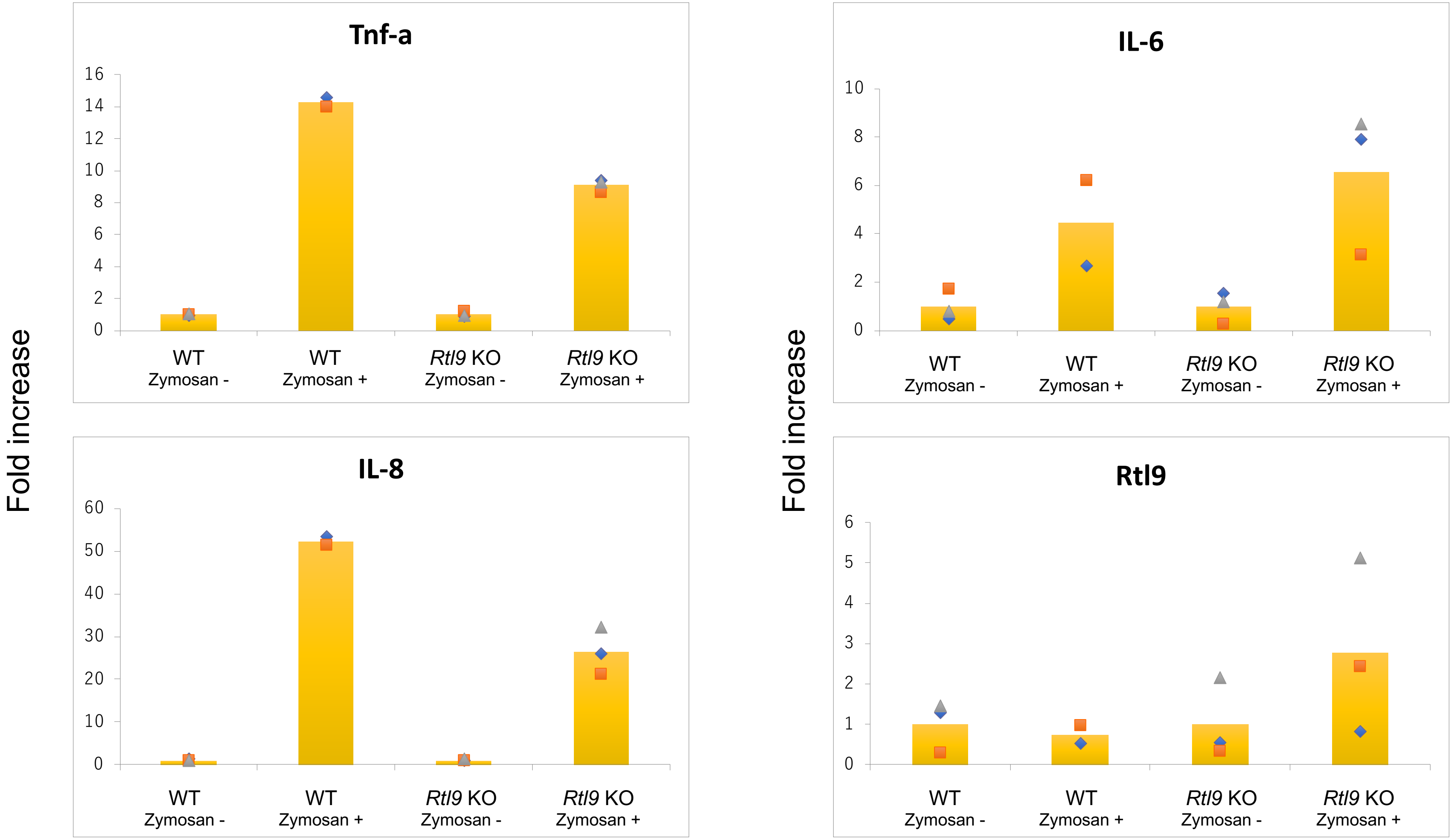


Fig. S13 Zymsan induces *Tnfa*, *Il6* and *Il8* expression in both WT and *Rtl9* KO microglia

### **Fig. S13 Zymosan induces *Tnfa*, *Il6* and *Il8* expression in both WT and *Rtl9* KO microglia**

*Tnfa, Il6, Il8* and *Rtl9* mRNA were analyzed at 0 min (no administration) and 30 min after zymosan administration to WT and *Rtl9* KO microglia (n=3 each). The expression levels are averaged with non-administration samples as 1. *Rtl6* primers were designed for the 3'-UTR, and were therefore also present in the *Rtl9* KO microglia. It should be noted that the *Rtl9* mRNA itself increases upon zymosan administration in *Rtl9* KO microglia, presumably due to feedback regulation of the RTL9 protein.

The PCR primers used were as follows:

*Tnfa,* 5′-GACAAGGCTGCCCCGACTACG-3′ (forward) and 5′-CTTGGGGCAGGGGCTCTTGAC-3′ (reverse); *Il6*, 5′-AGTTGCCTTCTTGGGACTGA-3′(forward) and 5′-CCTCCGACTTGTGAAGTGGT-3′ (reverse); *Il8*: 5′-CTCTCAAGGGCGGTCAAAAAGTT-3′ (forward) and 5′-TCAGACAGCGAGGCACATCAGGTA-3′(reverse); *Rtl9*: 5'-TCACCTACATGCCTGTGACC-3' (forward) and 5'-CAACAACACCACATTGTTACGG-3' (reverse).
